# Discrete protein condensation events govern calcium signal dynamics in T cells

**DOI:** 10.1101/2024.07.31.606035

**Authors:** Shumpei Morita, Mark K. O’Dair, Jay T. Groves

**Affiliations:** Department of Chemistry, University of California-Berkeley; Berkeley, CA 94720, USA

## Abstract

Calcium level variations, which occur downstream of T cell receptor (TCR) signaling, are an essential aspect of T cell antigen recognition. Although coordinated ion channel activities are known to drive calcium oscillations in other cell types, observations of nonperiodic and heterogeneous calcium patterns in T cells are inconsistent with this mechanism. Here, we track the complete ensemble of individual molecular peptide-major histocompatibility complex (pMHC) binding events to TCR, while simultaneously imaging LAT condensation events and calcium level. Individual LAT condensates induce a rapid and additive calcium response, which quickly attenuates upon condensate dissolution. No evidence of cooperativity between LAT condensates or oscillatory calcium response was detected. These results reveal stochastic LAT protein condensation events as a primary driver of calcium signal dynamics in T cells.

**One-Sentence Summary:** Ca^2+^ fluctuations in T cells reflect stochastic protein condensation events triggered by single molecular antigen-TCR binding.

## Main Text

Calcium elevation is a crucial step in T cell activation that regulates essential downstream pathways, including nuclear factor of activated T cells (NFAT) activation, cytotoxicity, and cytoskeleton reorganization (*1*–*3*). As is broadly observed in many cell types, intracellular calcium levels in T cells exhibit significant temporal variation upon stimulation, which is thought to encode information for downstream pathways (*4*). For example, apparent calcium oscillations (or fluctuations) in T cells have been suggested to regulate downstream transcription factors in a frequency-dependent manner (*5, 6*). Different temporal patterns affect cytoskeletal activity, regulating cell shape, polarity, motility (*7*–*10*), as well as CD69 upregulation and perforin secretion (*11, 12*). Optogenetic manipulation of calcium signals in T cells has further demonstrated the possibility of temporally encoded information modulating immunological outcome (*13*). The importance of calcium signal dynamics in regulating T cell behavior is broadly appreciated. However, comprehensive understanding of the underlying mechanisms is still lacking—a fact underscored by the wide variety of empirical parameterization and categorization methods employed for interpreting calcium measurements (*14*–*18*).

Calcium fluctuations in T cells significantly contrast several other cell types, such as hepatocytes, smooth muscle cells, or mammalian oocytes, which exhibit clearly periodic calcium oscillations. Oscillations in these cells are generated from feedback loops among signaling molecules and ion channels, mainly involving inositol 1,4,5-triphosphate (IP_3_) and IP_3_ receptor (IP3R), which induce the oscillatory efflux of calcium ions from endoplasmic reticulum (ER) stores (*19, 20*). In T cells, on the other hand, the primary source of cytosolic calcium elevation is influx from the extracellular space through the store-operated calcium entry (SOCE) pathway (*4,21,22*). Some early studies suggested a delayed negative feedback loop involving SOCE induction and ER calcium store refilling, along with related ion transport activities, could generate calcium oscillations in T cells (*23*–*30*). However, observed calcium dynamics in T cells markedly differ from the periodic oscillations predicted by this type of model. While some early studies of T cell calcium signaling reported apparently oscillating behavior (*4, 31*), it later became clear that T cells exhibit heterogeneous calcium patterns including non-periodic fluctuations, transient spiking, and sustained elevation (*14, 32*). Additionally, calcium responses in T cells are dependent on stimulation dose with transient spikes and fluctuating patterns observed at low to intermediate doses, while high doses tend to induce an initial rise followed by slow decay with minimal fluctuations (*8,33*–*35*). Although the SOCE pathway and ion channel activities possess the potential of inducing oscillatory output, these experimental discrepancies taken together motivate a more detailed analysis of the origins of calcium signal dynamics in T cells.

A distinctive feature of the TCR signaling pathway is its extreme sensitivity; a few individual agonist pMHC molecules are sufficient to fully activate a T cell (*36*–*40*). Under these low antigen conditions, individual pMHC:TCR binding events occur in isolation, creating a stochastic sequence of discrete TCR activation signals (*41*). TCR triggering leads to calcium influx through a multistep pathway involving nucleation of protein condensates of the membrane-associated scaffold protein, linker for activation of T cells (LAT) (**Fig. 1A**) (*42*). TCR triggering by pMHC induces a phosphorylation cascade involving the recruitment and activation of Zap70, which phosphorylates multiple tyrosine residues on LAT (*43*–*45*). LAT then recruits adaptor and signaling proteins including Grb2, SOS, Gads, SLP-76, and PLCγ1 into a crosslinked protein network (*46, 47*), which was later found to form discrete condensates through a type of phase transition (*48, 49*). PLCγ1 is activated in the LAT condensates (*50, 51*), and subsequently activates calcium ion efflux from the ER through IP_3_ production and IP_3_ receptor activation (*52*). This efflux readily depletes intracellular calcium stores and activates the SOCE pathway, in which Stim proteins activate Orai channels at the plasma membrane to allow extracellular calcium influx (*1,53*–*56*).

**Fig. 1.**
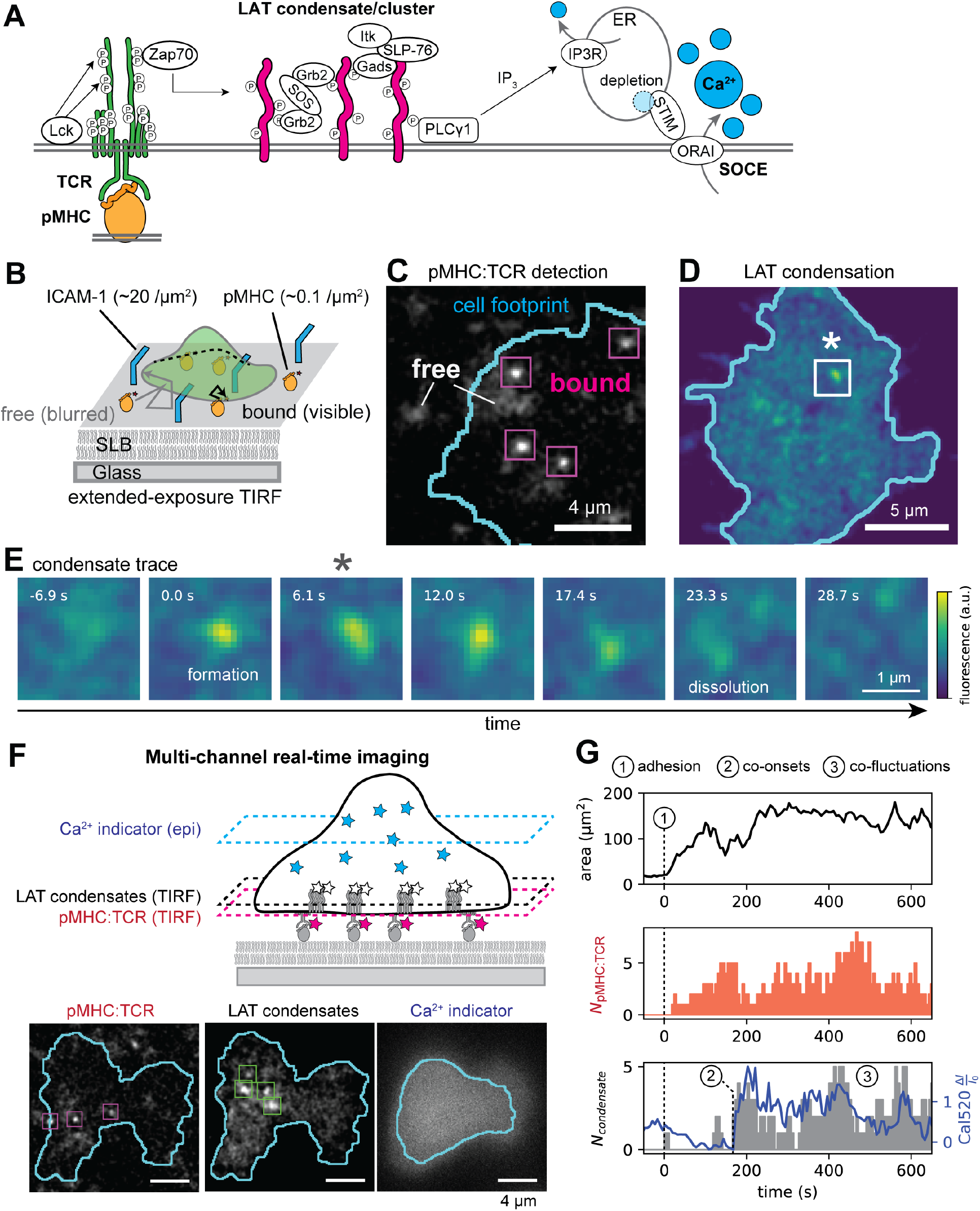
Observation of temporal correlation among pMHC:TCR binding, LAT condensates, and calcium levels. (**A**) Schematic of TCR signaling pathway. pMHC:TCR binding activates Lck and Zap70, which phosphorylates the tyrosine residues of LAT (shown as circled “P”). LAT condensate forms involving various proteins, including Grb2, SOS, Gads, SLP-76, Itk, PLCγ1. PLCγ1 activates IP3R through IP_3_ production, which activates SOCE pathway involving STIM and ORAI, leading to calcium ion influx. (**B**) Schematic of the pMHC:TCR complex imaging system. pMHC molecules bound to TCR on T cells exhibit lower mobility than free pMHC and are selectively imaged by extended-exposure TIRF microscopy. (**C**) Representative image of the selectively visualized bound pMHC molecules and blurred free molecules. (**D**) Representative TIRF image of LAT exhibiting discrete condensation. (**E**) Time series of a LAT condensate undergoing formation and dissolution, corresponding to a white box with asterisk in **C**. (**F**) Multi-channel imaging of pMHC:TCR complexes, LAT condensates, and Ca^2+^ indicator (Cal520). Snapshots of a representative cell at a representative frame are shown (bottom). (**G**) Representative single-cell traces of the cell-SLB interface area, the momentary number of pMHC:TCR (*N*_pMHC:TCR_), the momentary number of LAT condensates (*N*_condensate_) shown as grey bars, and the calcium level (Cal520 Δ*I* / *I*_0_) shown as blue line.

From the perspective of calcium signaling dynamics, the kinetic influences from upstream TCR signaling steps have received little attention. Existing calcium oscillation models implicitly assume that upstream TCR signaling is constant or well averaged over many molecular events. The effects of extremely low copy numbers of pMHC:TCR binding and LAT condensation events—and the intrinsic stochastic variation they inject into the system—have not been characterized. Here we describe experiments that directly track the kinetic connection between discrete LAT condensation events and calcium influx in a live, primary T cell system. We address two questions: *i*) if and when individual LAT condensates become functional to activate downstream calcium activity; and *ii*) if cellular level calcium fluctuations reflect these stochastic upstream signaling events.

Using a hybrid live cell-supported membrane system, we performed real-time imaging of single molecule pMHC:TCR binding events, LAT condensate formation and dissolution events, and the intracellular calcium level in living primary T cells. The momentary number of coexisting LAT condensates and the simultaneous calcium level exhibit a tight temporal correlation. Quantification of this response indicates that each LAT condensate contributes incrementally, and additively, to the cell-wide calcium level. No cooperativity among LAT condensates was observed; each condensate appears to be fully functional on its own within less than ∼10 sec of formation. Similarly, incremental reduction in intracellular calcium levels occurred on the same timescale upon LAT condensate dissolution, revealing that the calcium response is overdamped and does not exhibit any oscillatory or sustained activity. Lastly, spectral analyses of the LAT-calcium response reveal that the stochastic sequence of LAT condensate formation events is sufficient to describe the observed calcium fluctuations over the frequency range of 5-20 mHz. These results reveal that the discrete LAT condensation process in T cell signaling is a functional element gating signaling between TCR and calcium. Furthermore, stochasticity of the discrete LAT condensation events is a major contributor to the observed cell-wide calcium signaling dynamics.

### Simultaneous imaging of pMHC:TCR, LAT condensation, and calcium level

Primary AND-TCR T cells were imaged during interaction with supported lipid bilayers (SLB) functionalized with fluorescently labeled pMHC molecules (moth cytochrome C 88-103 peptide (MCC) with Atto647N) and ICAM-1 (∼20 molecules μm^-2^). Both proteins in the SLB are laterally mobile (*57, 58*) and the typical density of pMHC molecules (∼0.1 molecules μm^-2^) was low enough for well-resolved single molecule imaging. Upon contact with the SLB, ICAM-1 binding to LFA-1 on the T cell leads to adhesion and spreading to form a relatively uniform interface between the cell and the SLB, as observed by reflection interference contrast microscopy (RICM). Single pMHC molecules were imaged and tracked using total internal reflection fluorescence (TIRF) microscopy. Bound pMHC:TCR complexes were readily distinguished from free pMHC molecules based on mobility (**Fig. 1B, C**), as previously described (*41,58,59*). Individual pMHC molecules were observed to bind and unbind from TCR within this nascent immunological synapse; the distribution of binding dwell times is essentially exponential with mean around 50 sec, reflecting the strong pMHC:TCR interaction with MCC peptide (*41,49,58,59*). In occasional experiments where higher densities of pMHC (>0.5 molecule μm^-2^) were used, individual pMHC:TCR complexes were imaged in ensemble to confirm binding.

LAT and calcium levels were simultaneously observed, along with pMHC:TCR binding events, using T cells transduced with LAT-mCherry and loaded with the calcium-responsive fluorescent probe Cal520. As cells adhered to the SLB and began to experience pMHC:TCR binding events, discrete LAT condensates were observed to form and dissolve, as previously reported (*49*) (**Fig. 1D, E**). Epi-fluorescence imaging of the Cal520 signal was converted to the normalized difference from initial intensity (Δ*I* / *I*_0_) and used as a relative indicator of cytosolic calcium ion concentration. The dynamic range of the Cal520 response was calibrated using cells under calcium depletion and saturation conditions. The estimated dynamic range sufficiently covered the observed calcium levels in the present study, for which Δ*I* / *I*_0_ ranges approximately from 0 to 6 (**Fig. S1A-C**). Photobleaching of Cal520 was minor (about 10 to 20% bleached) and corrected using experimentally determined photobleaching curves (**Fig. S1D, E**).

Representative images of single-molecule pMHC:TCR binding events, LAT condensate formation, and calcium signal are depicted in **Figs. 1F** and **1G** (also see **Movie S1**). After cells land on the SLB, adhesion and spreading were observed as a quick increase in the interface area. We observed delays ranging from zero to 5 min between the initial cell-SLB contact and adhesive spreading. After T cell adhesion and spreading into a nascent immunological synapse, pMHC:TCR binding events began to occur. pMHC has a slow binding on-rate to TCR (in contrast to anti-TCR Fab’ or antibody (*60, 61*)), and individual pMHC:TCR binding events were well spread out in time. LAT condensation and calcium signals initially remained low. After delays ranging from zero to 150 sec, discrete LAT condensation events and calcium responses abruptly started. The momentary number of coexisting LAT condensates (*N*_condensate_) and the calcium level in each cell exhibited tightly correlated fluctuation patterns. We note that the Jurkat cell line exhibited altered behaviors and the results described here are distinct to primary T cells (**Supplementary Text S1**).

### Calcium levels exhibit rapid and additive response to individual LAT condensation events

The temporal patterns of *N*_condensate_ and the calcium level were highly similar within each cell, yet exhibited significant diversity among cells. Representative examples of simultaneous measurements of *N*_condensate_ and calcium level for individual cells at low pMHC density (0.10 ± 0.03 molecules μm^-2^) are plotted in **Fig. 2A**. Calcium level traces from some cells exhibited an abrupt onset and non-periodic fluctuations (cell 2, 3 in **Fig. 2A**), while in some cases, transient calcium spikes corresponding to individual LAT condensation events were clearly visible (cell 1). At higher pMHC densities (1.1 ± 0.2 molecules μm^-2^), calcium levels also tracked tightly with *N*_condensate_ in individual cells (**Fig. 2B**). Imaging variation due to changes in cell thickness were estimated using calcium-independent fluorophore Calcein and found to be relatively small, within 20% (**Fig. S3**). We note that the overall magnitude of apparent calcium levels varied among cells, even with similar levels of *N*_condensate_, part of which presumably stems from normalization inaccuracy due to the variation in the initial calcium levels. In the following data interpretations, we consider this intrinsic experimental limitation by averaging among multiple cells or focusing only on the relative fluctuations.

**Fig. 2.**
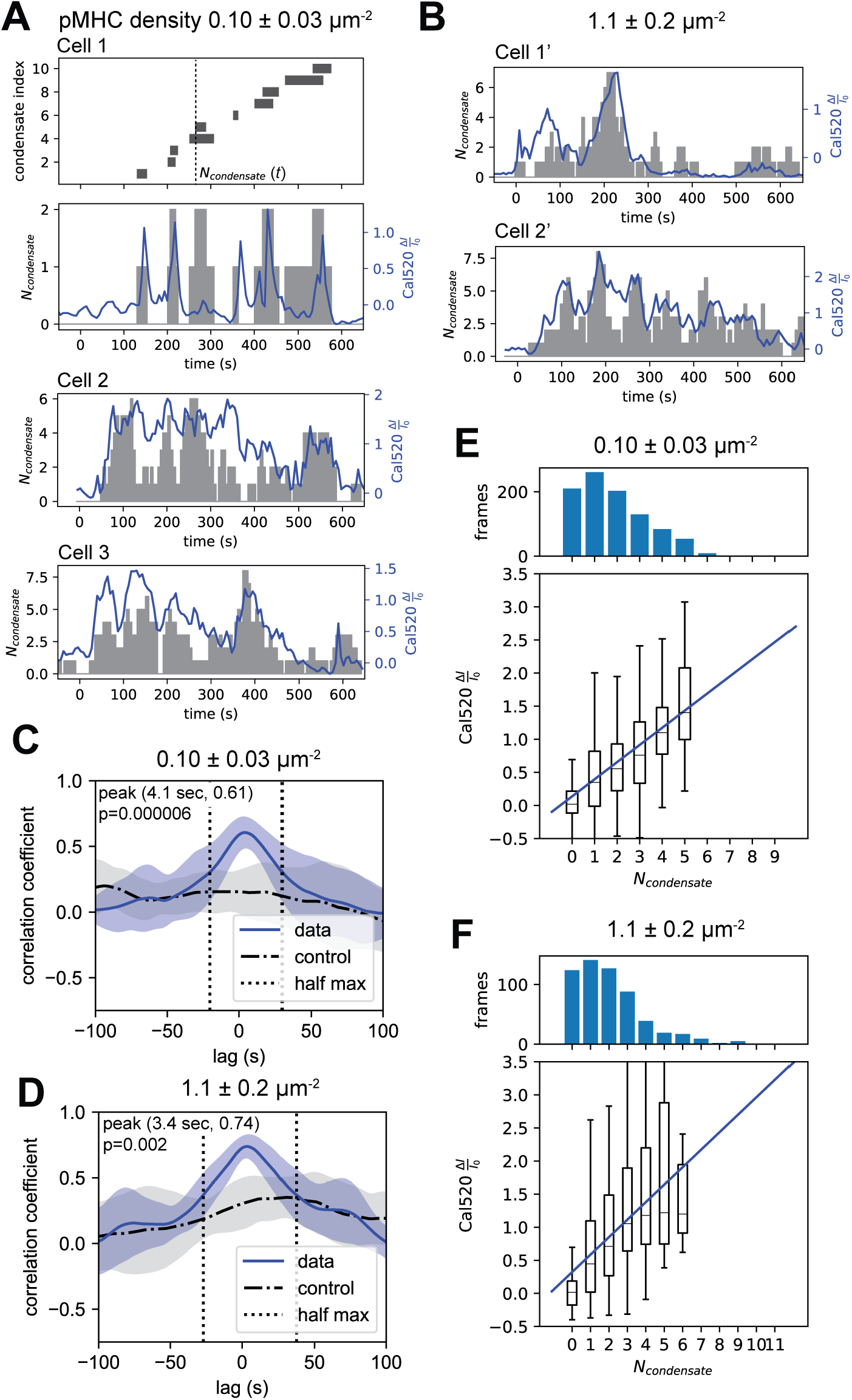
Co-fluctuating single-cell traces of LAT and calcium signals. (**A, B**) Representative traces of the momentary number of LAT condensates (*N*_condensate_, grey bars) and the calcium levels (Cal520 Δ*I* / *I*_0_, blue line) exhibiting the similarity within each cell and the diversity among cells. *N*_condensate_ was defined by counting the number of condensates co-existing at a given moment (cell 1, top), which correlated with calcium levels (cell 1, bottom). Additional representative cells are shown in **A** for lower and **B** for higher pMHC densities (density uncertainties indicate the min-max range used). (**C, D**) Cross-correlation between *N*_condensate_ and calcium traces was calculated for each cell. Mean ± SD and half-maximum lag values are shown. Cross-correlation between two traces from two different cells (cycled cell index) are shown as false-correlation control. p-values at the peak are shown (two-sided Welch’s t-test). (**E, F**) Response curves were constructed by frame-wise statistics, pooled from all cells. Frame-wise value-pairs (box plot) with linear regression (solid line) are shown. Data was collected from 10 cells from 3 mice for lower density, 6 cells from 3 mice for higher density.

Following the onset of LAT condensation events and calcium activity, a period of sustained and tightly correlated fluctuations in *N*_condensate_ and calcium level ensued. Cross-correlation analysis of *N*_conensate_ and calcium level traces revealed a peak with minimal lag (less than 4 sec) and narrow width (20 to 40 sec half width at half maximum), which shows that calcium level follows the same fluctuation pattern as *N*_condensate_ with no measurable delay and no trailing oscillatory or decay process (**Fig. 2C, D**). This non-hysteretic characteristic allowed measurement of the response function of the calcium levels to *N*_condensate_ (**Fig. 2E, F**). Calcium response is highly sensitive, clearly responding to even a single condensate, and additive, gradually increasing over the observed *N*_condensate_ range of 0 to 6. The response curves with different pMHC densities did not show noticeable difference, implying that the calcium response is primarily dependent on the copy number of discrete LAT condensates. The response was approximately linear in the unit of either Δ*I*/*I*_0_ or absolute concentration estimate (**Fig. 2E, F,S4A, B**). These response characteristics indicate that individual discrete LAT condensates each induce an essentially immediate calcium response, which quickly attenuates upon the dissolution of the condensate.

We confirmed that the observed calcium responses result from the calcium ion influx at the plasma membrane induced downstream of LAT condensation. During active pMHC:TCR-LAT-calcium signaling, LAT condensation was inhibited by the addition of PP2, a Src-family kinase inhibitor that inhibits Lck and other kinases (*62*). Immediately after the addition, LAT condensates stopped forming and calcium level dropped to baseline (**Fig. S5A**). While PP2 also inhibited the cell adhesion as previously reported (*62*), pMHC:TCR complexes were still existent at the cell-SLB interface, suggesting that the steps between TCR triggering and LAT condensate nucleation were inhibited by PP2 (**Fig. S5B**). On the other hand, the addition of EGTA to deplete extracellular calcium ion immediately quenched the intracellular calcium signal, while LAT condensate formation continued (**Fig. S5C, D**). These results confirm that calcium ion influx depends on LAT condensates, but LAT condensation does not depend on calcium signal.

### LAT condensation and calcium signals begin after a morphological kinapse-synapse transition

A distinct timepoint at which frequent LAT condensate formation and calcium level elevation initially appear was discernible in most of the cells (**Fig. 3A,S6**). Delay times between cell adhesion and the onset of LAT or calcium activity ranged from 0 to 150 sec for different cells, yet were simultaneous for each cell within the uncertainty of 25 seconds regardless of pMHC densities (**Fig. 3B, C**). We qualitatively observed that LAT-calcium signaling onset was accompanied by a morphological kinapse-synapse transition in some cells, which was indicated by the transition from asymmetric cell footprint with directional ameboid movement to the circular-symmetric footprint with centripetal membrane flow (**Fig. 3A,S6**) (*63*–*65*). These results suggest there exists an initial kinapse phase during which the first few pMHC:TCR binding events consistently fail to lead to LAT condensation and calcium signals, plausibly due to altered mechanical force or distinctly distributed signaling molecules (*65, 66*) (see **Supplementary Text S2** for extended discussion).

**Fig. 3.**
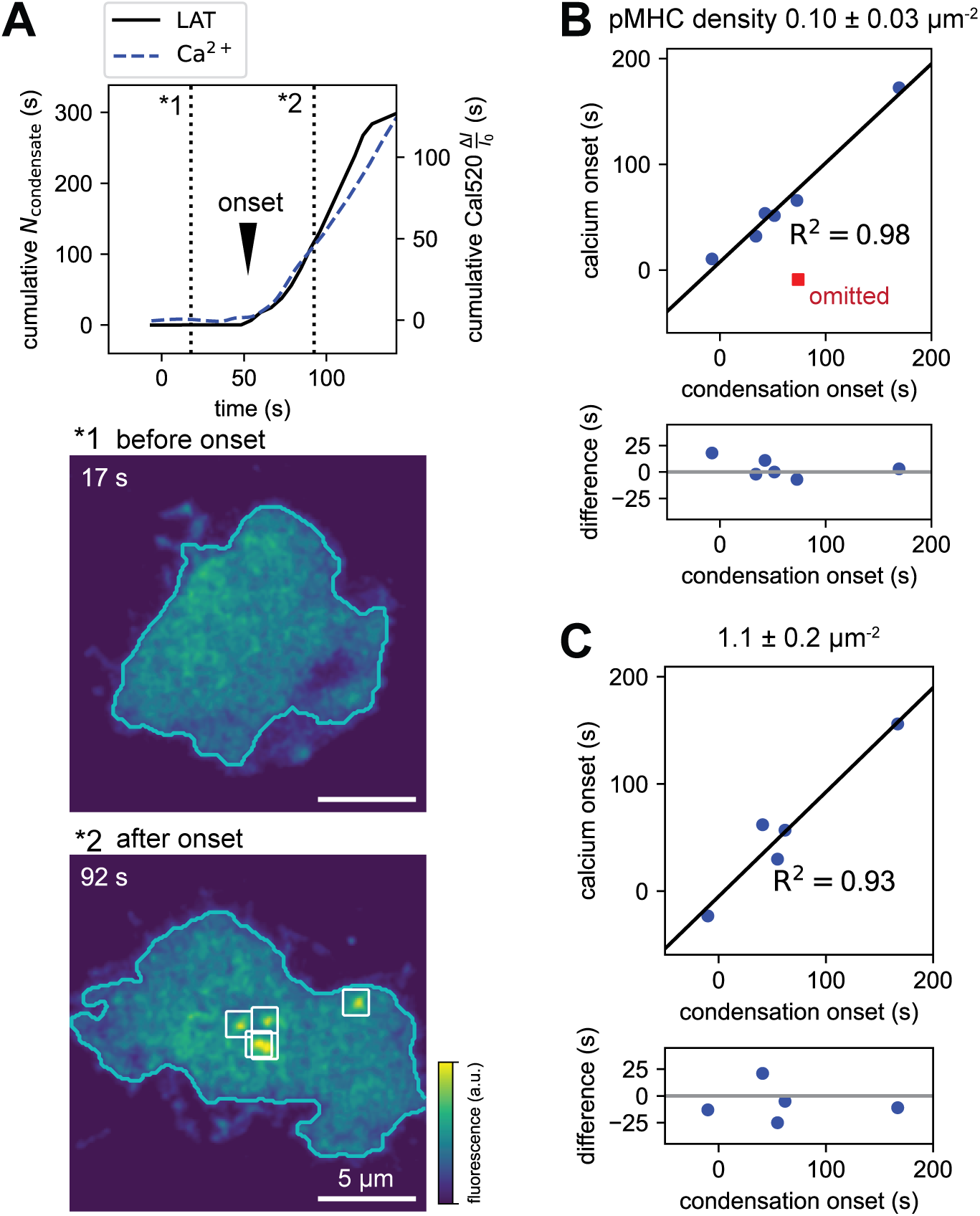
Simultaneous delayed onsets of LAT condensation and calcium signaling. (**A**, top) Representative cumulative traces of *N*_condensate_ and calcium signal for lower pMHC density. The onsets are identified as abrupt changes in the slopes (annotated with black arrow). (**A**, bottom) Representative LAT snapshots before (*1) and after (*2) the onsets. LAT condensates are annotated with white boxes. Morphological transition from asymmetric to symmetric cell footprint was observed. (**B, C**) Correlation analysis of the onsets of *N*_condensate_ and calcium signal for lower and higher pMHC densities. Pairs of onset delays from each cell are depicted with linear regression, exhibiting positive correlation (upper panels). The onset differences were about 25 sec or less (lower panels). 7/10 cells for lower density and 5/6 cells for higher density showed both detectable onsets. One data point with detection artifact was omitted from linear regression as an outlier (Grubbs test for the onset differences, α=0.05).

### Calcium signal dynamics change with increasing pMHC dose

We characterized the dose-dependence of calcium signaling in T cells over pMHC densities ranging from zero to 100 molecules/µm^2^ (**Fig. 4A**). These experiments were performed in non-transduced T cells to increase the experimental throughput and to provide corroborating control measurements without retroviral transduction. Cells adhering to SLB functionalized with only ICAM and without pMHC molecules typically exhibited transient calcium elevations with low overall intensity. As the pMHC density increased, observed calcium traces transitioned to spiking or fluctuating patterns (pMHC densities 0.12-0.21 μm^-2^), then to patterns with an initial rise followed by fluctuations (pMHC densities 0.53-4.6 μm^-2^). At even higher pMHC densities of 40-100 μm^-2^, calcium traces typically exhibited a strong initial rise followed by a slow decay with less fluctuating features. Calcium levels aligned well regardless of LAT-mCherry retroviral transduction (**Fig. 4B, C**).

**Fig. 4.**
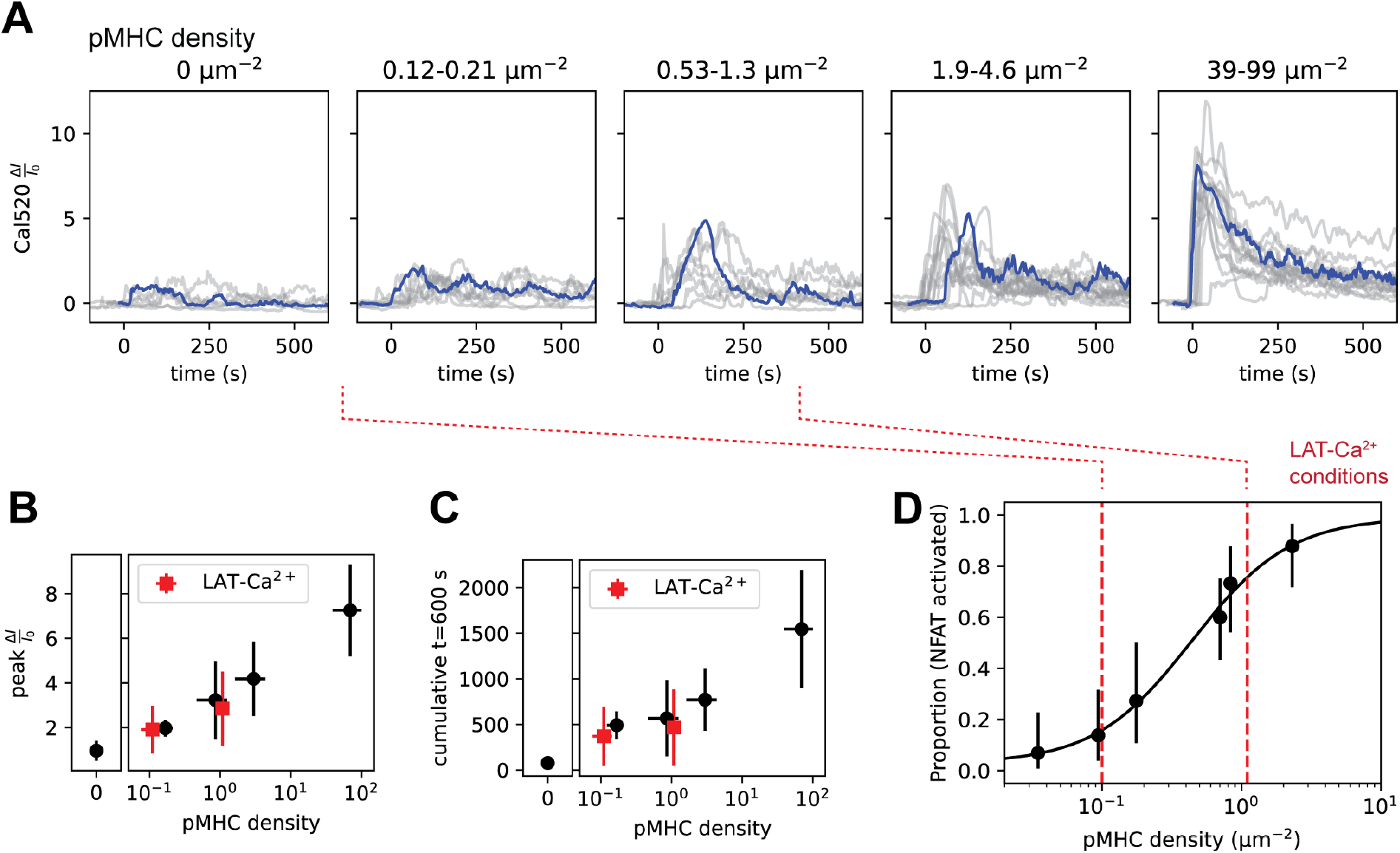
pMHC density-dependence of calcium response and NFAT activation. (**A**) Single-cell calcium traces from non-transduced T cells stimulated with different pMHC densities. Data was pooled from 3 mice, and each density range contains at least 7 cells from 2 mice. (**B, C**) Global quantification of calcium levels based on the peak height (**B**) and the cumulative intensity (**C**). The data from LAT-calcium simultaneous imaging experiments are overlayed in red, which reasonably aligned with non-transduced cell data. Error bars: SD (y-axis), min-max (density). (**D**) Representative titration curve of NFAT translocation assay. The proportions of the activated cells at each density are shown with 95% CI (Clopper-Pearson). Sigmoidal fit is shown as solid line. The pMHC densities with which LAT-calcium co-imaging was performed (red dashed lines) ranged over the NFAT activation threshold. A visual guide to show the corresponding densities in **A** is shown as red dotted lines.

We next calibrate the T cell functional response to this range of pMHC densities by tracking NFAT. NFAT translocation from cytosol to nucleus is a downstream event from intracellular calcium elevation and a marker of early T cell activation (*41,59,67,68*). For these experiments, T cells were transduced with an NFAT reporter consisting of regulatory domain of NFAT fused with mCherry (*68*). The proportion of the cells with nuclear-localized NFAT reporter is plotted against pMHC density, revealing an EC_50_ of 0.6 ± 0.2 pMHC molecules μm^-2^ (**Fig. 4D,S9**). The Cal520 treatment did not alter the NFAT activation threshold, validating it has minimal interference on calcium signaling (**Fig. S9**). The pMHC densities at which the LAT condensate-calcium relationship was characterized (0.10 ± 0.03 and 1.1 ± 0.2 μm^-2^) span over the NFAT activation threshold (0.6 μm^-2^), confirming that the calcium signals being studied are functional in T cell activation. The measured NFAT-activation threshold in these experiments approximates the proliferation threshold (0.2 μm^-2^) determined in a previous study (*69*) and the maximum pMHC densities tested here (40-100 μm^-2^) correspond to the densities used in previous studies to stimulate T cells thoroughly (ranging from 60 to 200 μm^-2^) (*62, 69*).

A widely adopted T cell stimulation strategy utilizes glass-immobilized anti-TCR:CD3. We tested this method and observed that glass surfaces coated with high-density anti-TCR induce qualitatively different behaviors in LAT condensation and calcium response. Under these conditions, LAT immediately condensed throughout the interface and dissolved quickly in about 1 min. The corresponding calcium levels showed an immediate rise followed by a slow decay (**Fig. S10A, B**), which is consistent with a previous study (*70*). At 1000-fold lower anti-TCR levels, some cells exhibited discrete LAT condensation and calcium fluctuations resembling behaviors observed with pMHC (see **Supplementary Text S3, Fig. S10C-E**).

### Discrete LAT condensates originate both proximally and distally to pMHC:TCR

Some of LAT condensates form in the vicinity of an individual pMHC:TCR complex (*49*). In the present experiments, these pMHC:TCR-proximal condensates accounted for less than about 20% of the total observed (**Fig. S11; Supplementary Text S4**). We investigate two plausible origins of the LAT condensates without colocalized pMHC (here referred to as pMHC:TCR-distal LAT condensates). First, Zap70 might contribute to distal LAT condensation through release from pMHC:TCR (*71*), TCR that remained active after pMHC unbinding (*72, 73*), or LFA-1-dependent activity (*74*). We observed that Zap70 was detectable in 66% of proximal and 57% of distal LAT condensates, confirming Zap70-LAT association commonly persists distal to pMHC:TCR (**Fig. S13**). Secondly, LAT can be trafficked to the plasma membrane in vesicular compartments, characterized by a SNARE protein Vamp7 (*75*–*78*). Imaging Vamp7 along with pMHC:TCR and LAT revealed 20% of apparent LAT condensates also contain Vamp7 (**Fig. S14**). Both Zap70 and Vamp7 signals were weak, thus their detection levels represent lower bounds on inclusion rate. The strong correlation between total LAT condensate count and calcium levels that we observed indicates that both proximal and distal condensates contribute to the cellular calcium signal dynamics (see **Supplementary Text S5** for extended discussion).

### The stochastic time sequence of LAT condensation events is sufficient to describe the calcium fluctuation behavior

As a minimal hypothesis, each LAT condensate formation event might simply trigger a pulse of calcium signal, independent of condensate lifetime or size, which sums to drive the fluctuating calcium level. We tested this minimal interpretation from a systems analysis perspective using a linear time-invariant (LTI) approximation. Condensate formation events were observed to occur at a near-constant rate, approximating a Poisson process (**Fig. 5A**). In the following analysis, these condensation events (input) produce pulses of calcium signal (output) via a transformation process described by an impulse-response function. This impulse-response function is a measurement of any oscillatory or sustained calcium response to a signal input for LTI systems. Data from all cells were pooled to determine the empirical impulse response function for calcium signaling in T cells (**Fig. 5B,S16**), which exhibits a singular peak with approximately 15 second delay and 20 second width. Consistent with the measured cross-correlation (**Fig. 2C, D**), no evidence of oscillation was detected. Convolution of this impulse response function with the experimental LAT condensate formation time sequence successfully reproduced the observed calcium traces (**Fig. 5C,S17, S18**). For comparison, we examined two other minimal models adding condensate lifetime (input of *N*_condensate_) and condensate size (input of the summed momentary condensate size Σ*I*_*condensate*_) onto the formation time sequence. Inclusion of this additional information did not improve the model fit to data, suggesting that the condensate formation time sequence alone is sufficient to describe the overall calcium behavior (**Fig. S17**). Furthermore, the experimental power spectral density (PSD) of calcium fluctuations matched the simulated fluctuation levels for the formation time sequence following a Possion process over the frequency of about 5 to 20 mHz (period of 50 to 200 sec) (**Fig. 5D,E, S18**). Notably, this frequency range matches reports of apparent oscillations in previous studies, typically with period of 80 to 140 sec (*4,21,31,35*). These results confirm that a major portion of the measured calcium signal is fully described by a linear transformation of the LAT condensation event sequence, without underlying oscillatory or feedback circuits (see **Supplementary Text S6** for complete descriptions).

**Fig. 5.**
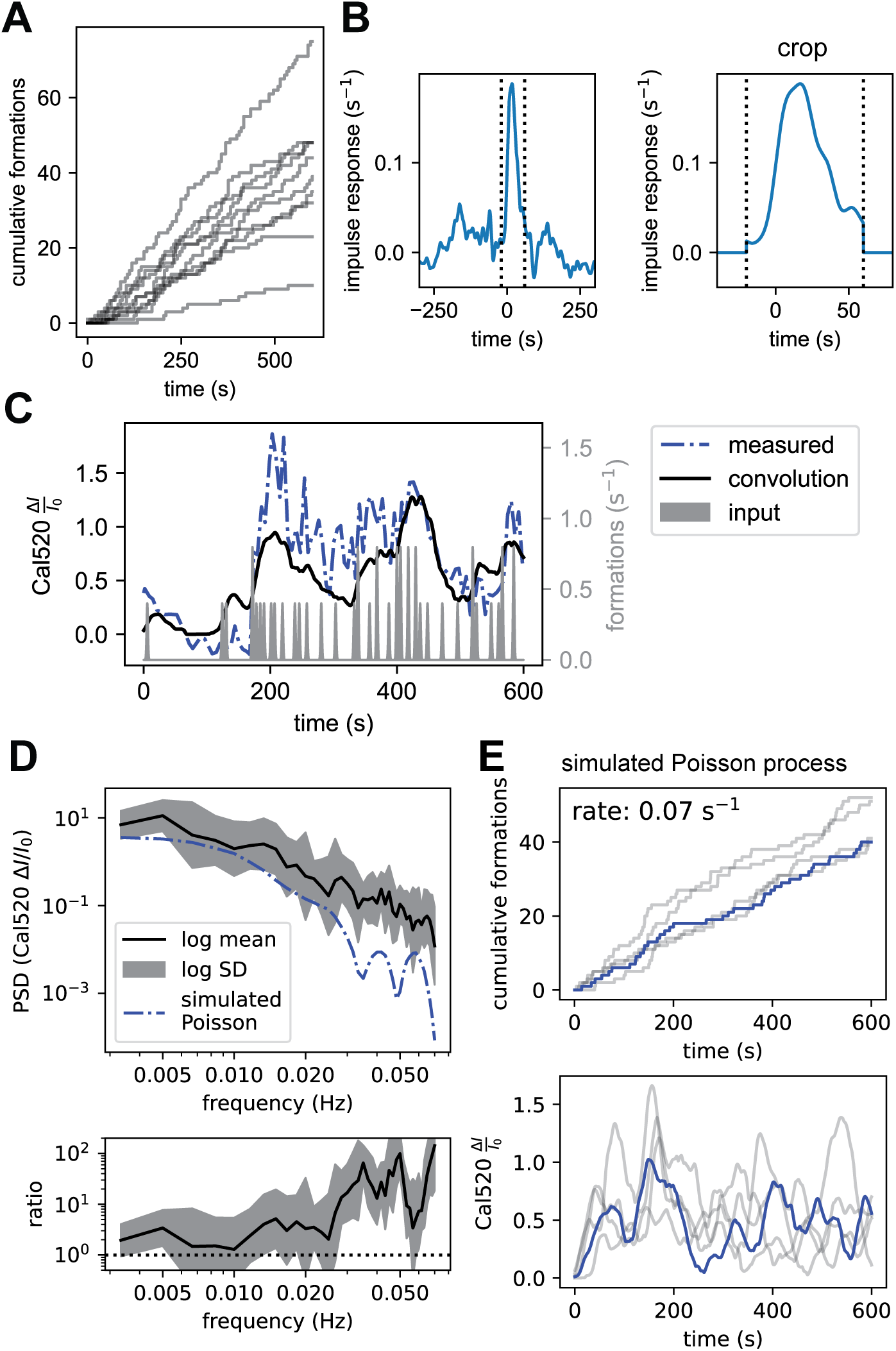
Stochastic LAT condensate formation events with calcium signal pulses recapitulate observed fluctuations. LTI system analysis for lower pMHC densities (0.10 ± 0.03 µm^-2^) is shown. (**A**) Cumulative traces of condensate formation events from each cell. (**B**) Empirical impulse response was determined by considering the condensate formation events as the input (expressed as gaussian spikes) and the calcium trace as the output. The impulse response was cropped to extract the main peak (right panel). (**C**) Representative cell traces of the formation events (input) and the measured calcium level. The convolution of the impulse response and the input is overlaid, recapitulating the measured calcium trace. (**D**) PSD of calcium traces (logarithmic mean ± SD) is compared with simulated Poisson noise (top). The ratio of experimental PSD and simulated PSD is also shown (bottom). (**E**) Representative simulated traces of condensate formation events following Poisson process with rate 0.07 s^-1^ (top) and the convoluted calcium traces (bottom). Representative pair of traces is highlighted in blue.

## Conclusion

Overall, the experiments described here reveal several distinct features of the role of LAT condensation in T cell signaling. At antigen densities near the activation threshold for T cells, LAT condensates remain discrete and generally well separated from one another. We find that each LAT condensate becomes fully active with respect to calcium signaling quickly after nucleation, and no evidence for cooperativity between condensates is measurable. This suggests rapid and independent signal propagation from PLCγ1 activation in the LAT condensate to SOCE induction. Each condensate makes an additive contribution to the cellular calcium level and we observe no oscillating or trailing responses. The calcium signaling system in T cells is apparently overdamped and differs from the clearly oscillatory behaviors that have been observed in other cell types. Rapid calcium attenuation has also been reported upon a sudden shut-off of TCR stimulation using optogenetic methods (*79*), which is consistent with the overdamped system response measured here. We confirmed that calcium responses corresponding to individual LAT condensates originate from extracellular calcium ion influx and not efflux from ER. This observation aligns with recent reports that even early or transient calcium signals with or without TCR stimulation are SOCE-dependent (*8, 80*). Lastly, experimental response characteristics as well as spectral analyses indicate that cellular calcium fluctuations are largely reflective of underlying stochastic LAT condensation events, and ultimately the individual pMHC:TCR molecular binding events that initiate them.

## Acknowledgments

We thank F. Marangoni (Harvard Medical School) for providing the NFAT reporter plasmid; L. Teyton (Scripps Research Institute) and M. Davis (Stanford University) for providing the MHC and ICAM-1 bacmids.

## Author contributions

Conceptualization: JTG, SM

Methodology: SM, MKO

Investigation: SM

Supervision: JTG

Writing – original draft: SM, JTG

Writing – review & editing: SM, JTG, MKO

Funding acquisition: JTG, SM

## Data and materials availability

The data and materials newly created in this study are available from the corresponding author (JTG) upon reasonable request.

## Supplementary Materials

Materials and Methods

Supplementary Text S1 to S6

Figs. S1 to S26

Table S1 Movie S1

References (*81*-*97*)

