## supplementary_information for "Discrete protein condensation events govern calcium signal dynamics in T cells"

### Discrete LAT protein condensation and dissolution events govern $Ca^{2+}$ signal dynamics in T cells

**The PDF file includes:**

Materials and Methods
Supplementary Text S1 to S6
Figs. S1 to S26
Table S1
References (*81*-*97*)

**Other Supplementary Materials for this manuscript include the following:**

Movie S1

### Materials and Methods

Supported lipid bilayer and imaging chamber preparation

Mouse or human ICAM-1 extracellular domain with His-10 tag and MHC class II I-$E^k$ with two His-6 tags were expressed and purified as previously described (*57*). Peptide-loaded MHC molecules were prepared essentially as previously described (*58*). MCC peptide (ANERADLIAYLKQATK) and MCC-GGSC peptide (ANERADLIAYLKQATKGGSC) were synthesized on campus (D. King, Howard Hughes Medical Institute Mass Spectrometry Laboratory at University of California, Berkeley) or commercially (Elim Biopharmaceuticals, Hayward, CA). MCC-GGSC was labeled with Atto647N-maleimide (ATTO-TEC), purified by C18 reversed phase column HPLC, and identified by MALDI-TOF mass spectrometry. Excess amount of MCC-GGSC-Atto647N (final concentration 20 μM) were incubated with MHC molecule (final concentration 1.8 μg/mL, 27 nM) in loading buffer (PBS acidified with citric acid to pH 4.5, 1% BSA) over night at 37°C. The mixture was diluted-concentrated with 10 kDa MWCO filters (Spin-X UF, Corning) using TBS twice to remove excess peptides, then used for bilayer functionalization. For the pMHC with 9.1% labeling fraction, non-labeled MCC peptide was also loaded to MHC molecule separately, which was mixed with the labeled MCC-GGSC-Atto647N loading mixture at 10:1 molar ratio before the dilution-concentration procedure. This partially labeled pMHC was used for SLB functionalization with higher densities than ~5 $\mu m^{-2}$.

Fab' ligand for Jurkat experiments was prepared following the methods used for other Fab' ligands (*61*, *81*). Single-strand DNA with thiol group at 3'-end and amino group at 5'-end (5'-/5AmMC6/GGT GTG ATG TAT GTG GA/3ThioMC3-D/ -3', Integrated DNA Technologies) was conjugated with Atto647N-maleimide (ATTO-TEC) and then SM(PEG)6 (Thermo Scientific) as described in these reports. Mouse anti-human CD3ε IgG (clone OKT3, Bio X Cell Catalog #BE0001-2, RRID: AB_1107632) was dialyzed to 0.1 M sodium acetate buffer pH4.1 at 4 °C to yield 1.1 mg/mL solution. IgG was digested by pepsin (IgG 1.0 mg/mL, pepsin 130 μg/mL) at 37 °C for 2 hours with occasional mixing. The digestion was quenched by adding 10 vol% of 1 M Tris-HCl buffer, pH8.5. White precipitate formed during digestion but dissolved upon neutralization. The buffer was then exchanged to PBS using 50 kDa MWCO filters (Amicon, Millipore-Sigma). By-product fragments were adsorbed by Pierce protein A agarose beads (Thermo Scientific) for 30 min at room temperature, then the mixture was centrifuged and the supernatant was collected. The obtained $F(ab')_2$ fragment was reduced by 2 mM 2-mercaptoethylamine in PBS with 2 mM EDTA at room temperature for 90 min. The buffer was exchanged to PBS with 2 mM EDTA using 30 kDa MWCO filters. The Fab' fragment (1 mg/mL) was conjugated to A647N-labeled single-strand DNA with maleimide group (3 to 4 molar ratio) by incubation at 4 °C over-night, then buffer-exchanged to PBS using 30 kDa MWCO filters. The conjugate was purified by size-exclusion chromatography (superdex S75 increase column with AKTA FPLC, Cytiva) and anion-exchange chromatography (Mono Q column, Cytiva). The reactions were tracked by non-reducing SDS-PAGE.

Small unilamellar vesicles (SUV) with 98 mol% 1,2-dioleoyl-sn-glycero-3-phosphocholine (DOPC, Avanti Polar Lipids) and 2 mol% 1,2-dioleoyl-sn-glycero-3-[(N-(5-amino-1-carboxypentyl)iminodiacetic acid)succinyl] nickel salt (DGS-NTA, Avanti Polar Lipids) was prepared by tip-sonication of 0.5 mg/mL lipid suspension in water followed by centrifugation (21000 G, 30 min, 4°C) to collect the supernatant. Then, supported lipid bilayer (SLB) was formed onto #1.5 glass coverslip set into Attofluor cell chamber (Invitrogen, Thermo Fisher). Coverslips were cleaned by sonicating in 1:1 water:2-propanol mixture, etched with piranha solution (1:3 mixture of 30% $H_2O_2$ and sulfuric acid), then rinsed by water and set into clean chambers. SLB was formed by adding 1:1 mixture of SUV and TBS into chambers and incubating for more than 30 min. The chambers with SLB were rinsed by TBS, then incubated with 10 mM $NiCl_2$ in TBS for 5 min to ensure the full activity of DGS-NTA. Imaging buffer was prepared with the following composition: 20 mM HEPES, 137 mM NaCl, 5 mM KCl, 1 mM $MgCl_2$, 1.8 mM $CaCl_2$, 0.1% w/v D-glucose, 0.1% w/v BSA, pH 7.2-7.4. The chambers were then rinsed by imaging buffer and incubated for more than 30 min to block the bilayer defects by BSA contained in the buffer, then used for functionalization. ICAM-1 (2 nM) and pMHC (concentration adjusted for each preparation based on the resulting density, estimated to be about 1-10 pM for 0.1-1 $\mu m^{-2}$) were added to the chambers and incubated for 30-40 min, then rinsed by imaging buffer. The ICAM-1 density is estimated to be ~20 molecule/$\mu m^2$ (*49*, *57*).

For Jurkat cell experiments, chamber preparation was performed essentially as described previously (*61*). SUV and SLB were prepared with 95 mol% DOPC, 2 mol% DGS-NTA, and 3 mol% 1,2-dioleoyl-sn-glycero-3-phosphoethanolamine-N-[4-(p-maleimidomethyl)cyclohexane-carboxamide] (sodium salt) (18:1 PE MCC). The maleimide groups in the bilayer was conjugated with thiol-functionalized single-strand DNA complementary to the Fab' ligand (5'-/5ThioMC6-D/CCA CAT ACA TCA CAC C -3', Integrated DNA Technologies). SLB was then functionalized with human ICAM-1 (40 nM) and OKT3 Fab' ligand (concentration adjusted for each preparation based on the resulting densities).

The chambers with glass-immobilized antibody were prepared essentially as previously described (*70*). The glass coverslips were cleaned and etched in the same way as the SLB chambers, set into the chambers, then incubated with the PLL in water (0.1 mg/mL, Sigma Aldrich) at room temperature for 15 min. The PLL solution was aspirated, then the chambers were dried in vacuum and stored at room temperature until used. The chambers were incubated with anti-mouse TCRβ IgG (clone H57-597, 10 μg/mL or 10 ng/mL, Bio X Cell Catalog #BE0102, RRID: AB_10950158) in PBS for 2 hours in 37 °C 5% $CO_2$ incubator, washed by PBS, and imaging buffer was added for imaging.

<u>DNA constructs</u>

Retroviral transduction was performed with murine stem cell virus (MSCV) vector with corresponding inserts. NFAT C2 (reg)-mCherry-P2A-LAT-EGFP constructs were prepared as previously reported (*49*). Other constructs were prepared by standard restriction enzyme cloning or Gibson cloning. LAT, NFAT, Zap70, Vamp7 sequences encode *Mus musculus* genes.

Lenti-viral transduction of Jurkat cells was performed using pHIV vector. Human LAT sequence fused with mCherry was cloned into pHIV vector by standard restriction enzyme cloning or Gibson cloning. Two helper plasmids pMD2.G (Addgene No. 12259) and psPAX2 (Addgene No. 12260) were used for packaging.

<u>Primary T cell preparation</u>

T cell culture medium was prepared with the following composition: DMEM with GlutaMAX (Gibco, Thermo Fisher) supplemented with 10% FBS, 1 mM sodium pyruvate, 2 mM L-glutamine, 1x Corning MEM nonessential amino acids (Fisher Scientific), 1x Corning MEM vitamins (Fisher Scientific), 0.67 mM L-arginine, 0.27 mM L-asparagine, 14 μM folic acid, 1x Corning Penicillin/Streptomycin (100 IU, 0.1 mg/mL respectively, Fisher Scientific), 50 μM β-mercaptoethanol. Primary AND-TCR T cells were prepared and cultured essentially as described previously (*82*). T cells were harvested from the lymph nodes and the spleen from F1 mice of B10.Cg-Tg(TcrAND)53Hed/J and B10.BR-H2k2 H2-T18a/SgSnJ (Jackson Laboratory, stock number 674005) and resuspended in T cell culture medium at ~10 M cells/mL (Day 1). Both male and female mice were used. To the cell culture was added 2 μM MCC peptide (ANERADLIAYLKQATK) at Day 1 for T cell priming by co-harvested splenocytes. IL-2 (25 ng/mL, Sigma-Aldrich) was added at Day 2 and the cells were kept under IL-2 stimulation with this concentration after this point. After Day 3, the cells were maintained in T cell culture medium with IL-2 at ~2.5 M cells/mL. The retroviral vector was produced with the corresponding genes at Day 3 or 4 using Platinum-E packaging system (*83*). The pMSCV plasmid was transfected to Platinum-E cells (Cell Biolabs, catalog #RV-101) with polyethylenimine 2 days before collecting viral supernatant. For transduction, the cells were mixed with freshly collected viral supernatant in T cell culture medium supplemented with IL-2 and polybrene (4 μg/mL, Sigma-Aldrich) and centrifuged (1500 G, 60 min, room temperature). The centrifuged cells were cultured untouched for one day and then resuspended in fresh media. The transduced cells were used 2 or 3 days after the transduction (Day 5-7) for imaging. Non-transduced cells were maintained in IL-2-containing media after priming and used at Day 5-7. All animal work was performed with prior approval by Lawrence Berkeley National Laboratory Animal Welfare and Research Committee under the approved protocols 177003 and 177005.

Jurkat cell preparation

Lentivirus for the transduction of human LAT-mCherry was packaged in HEK293FT cells (RRID: CVCL_6911). HEK293FT cells in DMEM supplemented with 10% FBS without antibiotics were transfected with pHIV-LAT-mCherry along with helper plasmids pMD2.G and psPAX2 using Lipofectamine LTX (Thermo Fisher). Viral supernatant was collected two and three days after transfection and concentrated using Lenti-X Concentrator (Takara Bio). Concentrated viral suspension was frozen in liquid nitrogen and stored at -80 °C.

Jurkat cells (from Cell Culture Facility, UC Berkeley) were maintained in RPMI medium supplemented with 10% FBS and 1x Corning Penicillin/Streptomycin. For imaging, cells were mixed with the viral suspension and polybrene (final concentration 10 μg/mL, Sigma-Aldrich) at the cell density of 0.5 M/mL. The virus-containing medium was replaced with fresh media after one day, and the cells were used for imaging three days after lentivirus addition.

Drug treatments for imaging experiments

For Cal520 loading, 2x working solution containing 5 μM Cal520-AM (AAT Bioquest) and 0.04% Pluronic F-127 (Invitrogen) in the loading buffer (20 mM HEPES, 137 mM NaCl, 1.8 mM $CaCl_2$, 5 mM KCl, 1 mM $MgCl_2$, pH 7.2-7.4, containing 0.1 % DMSO from Cal520-AM stock solution) was prepared, and mixed with 2.5 M cell/mL cell suspension in the culture medium with 1:1 volume ratio. After 20 min incubation in cell incubator (37 °C, 5% $CO_2$), the cells were centrifuged and re-suspended in fresh media (1.25 M cell/mL) and incubated for at least 10 min in the cell incubator to ensure the AM ester cleavage. The cells were rinsed with the imaging buffer and used for imaging within about 1 hour after dye loading. The non-responsive dye Calcein-AM was loaded similarly. 2x working solution containing 0.2 μM Calcein-AM (Cayman Chemical) in the loading buffer was prepared and mixed with 2.5 M cell/mL cell suspension in the culture medium with 1:1 volume ratio, incubated for 10 min in cell incubator. The cells were incubated in fresh media for at least 10 min, then rinsed by imaging buffer before used for imaging.

For the calcium depletion condition to validate the dynamic range of Cal520, cells were pre-treated with BAPTA-AM. The cells after Cal520 loading were precipitated and re-suspended in the culture medium supplemented with 20 μM BAPTA-AM (Cayman Chemical), 0.02% Pluronic F-127, 3 mM EGTA at the cell density of 2.5 M cell/mL, incubated for 20 min in cell incubator, then rinsed with the imaging buffer without $CaCl_2$. The cells were imaged in the imaging buffer without $CaCl_2$ supplemented with 1 mM EGTA. Under this calcium depletion condition the cells did not adhere well to the bilayer, but the epi-fluorescence imaging of Cal520 could still be performed. For the calcium saturation condition, ionomycin at high concentration (final concentration 13.38 μM, Sigma-Aldrich) and additional $CaCl_2$ (final concentration 10 mM) were added to the imaging chamber during the image acquisitions. The cells underwent calcium-induced necrosis several minutes after the addition. The imaging was performed quickly before the necrosis.

For the PP2 treatment, PP2 (final concentration 20 μM, 0.1% DMSO, Cayman Chemical) was added to the imaging chamber during the image acquisitions. For the extracellular calcium chelation experiments, EGTA (final concentration 3.6 mM or 6.2 mM) was added to the imaging chamber during the image acquisitions. More than two times higher EGTA concentration than the total $Ca^{2+}$ in the imaging buffer ensures the low concentration of free $Ca^{2+}$ in the scale of 10-100 nM.

For Jurkat cells, cells were first loaded with Cal520-AM following the same procedures as primary T cells. Then the cells were stained with DiD (Abcam). DiD solution in ethanol (100 μM) was diluted in PBS by 100-fold (1 μM final concentration), then bath-sonicated for 15 min to ensure sufficient dye suspension. The solution was mixed with cell suspension (1.25 M/mL) at 1:1 volume ratio, and the mixture was incubated for 5 min. The cells were rinsed by imaging buffer before imaging. For Jurkat cells, the imaging buffer contained 1 mM $CaCl_2$ to match the $Ca^{2+}$ concentration to RPMI medium. The imaging of Fab' ligands were performed with the cells treated with Cal520-AM and without DiD.

Microscope

TIRF microscopy was performed on a motorized inverted microscope (Nikon Eclipse Ti-E; Technical Instruments, Burlingame, CA) with Lumen Dynamics X-Cite 120 LED Fluorescence Illumination System (Excelitas Technologies, Waltham, MA, USA) and a motorized stage (MS-2000; Applied Scientific Instrumentation, Eugene, OR). A laser launch with 488, 561, and 637 nm diode lasers (Coherent OBIS, Santa Clara, CA) was aligned into a custom-built fiber launch (Solamere Technology Group Inc., Salt Lake City, UT). For TIRF imaging, laser excitation was illuminated through a four bands beam splitter (ZT405/488/561/640rpc) to the objective lens (NA 1.49, 100×, oil immersion, Apochromat TIRF, Nikon), then the emission light was filtered through an emission filter (ET525/50M, ET600/50M, or ET700/75M). For epi-fluorescence imaging, LED excitation light was illuminated through an excitation filter (ET470/40 or ET545/30) and a dichroic mirror (ZT488rdc or ZT561rdc), and the emission was collected through emission filter (ET525/50M or ET600/50M). For RICM, LED excitation light was illuminated through D546/10x excitation filter and 50/50 beam splitter. All optical filters were purchased from Chroma Technology Corp. (Bellows Falls, VT). Multi-channel imaging was performed with sequential switching among channels. The tube lens was chosen from two different magnification (1x and 1.5x). Emission was captured on EM-CCD (iXon Ultra 897; Andor Inc., South Windsor, CT). The sample and the objective lens were kept at 37 °C with temperature control system (CU-109, Live Cell Instrument). The equipment was controlled using the software MicroManager (*84*). The pixel size was 0.160 μm (1x tube lens) or 0.106 μm (1.5x tube lens) and the field of view was 512×512 pixels, 81.9 μm (1x tube lens) or 54.6 μm (1.5x tube lens).

Imaging

Imaging chambers with supported lipid bilayer (98% DOPC, 2% NTA-DOGS) functionalized with ICAM-1 (~20 molecules $\mu m^{-2}$) and Atto647N-labeled MCC pMHC filled with imaging buffer were prepared. The pMHC densities were determined by single-molecule imaging and intensity extrapolation prior to cell imaging (described below). Primary murine AND-TCR T cell blasts with proteins of interest retrovirally transduced were prepared. For the imaging of the cells just landing onto the bilayer, the cells pretreated with fluorophores or drugs were re-suspended in imaging buffer and small volume was added to the imaging chamber (about 1.5 μL of 6 M cell/mL, 37 °C on the heating stage) at the position above the objective lens, and the image acquisitions were quickly started. For the imaging of the cells after adhesion, larger volume of dilute cell suspension (about 100 μL of 2.5 M/mL) was added to the entire chamber.

For the measurement of LAT condensation and calcium levels, cells transduced with LAT-mCherry and loaded with Cal520 were imaged with 640 nm excitation TIRF (exposure time 500 ms, laser power density 3.3 $W/cm^2$), 561 nm excitation TIRF (100 ms, 2.1-3.6 $W/cm^2$), and 488 nm excitation epi-fluorescence (20 ms, focal plane height 4 μm) channels with 1.5x magnification tube lens, average frame interval of 6.3 sec. To capture the cells prior to adhesive spreading, LAT-mCherry-positive cells were quickly found using 561 nm epi-fluorescence channel (about 10 snapshots or less to minimize photo-toxicity). Cell adhesion, pMHC:TCR complexes, LAT condensates, and calcium levels were quantified and used for the following analyses.

For the measurements of calcium levels with different pMHC densities, non-transduced cells with Cal520 were imaged with RICM (20-50 ms) and 488 nm excitation epi-fluorescence (20 ms, 4 μm) channels with 1x magnification tube lens, average frame interval of 3.6 sec. For the densities higher than about 5 molecules $\mu m^{-2}$, partially labeled pMHC (9.1% labeled by fluorophore) was used.

NFAT translocation assay was performed using the regulatory domain of NFAT C2 fused with a fluorescent protein, which visualizes NFAT localization without interfering with transcriptions (*68*). The assay was performed essentially as previously described with minor modifications (*41*). Cells transduced with NFAT C2 (reg)-mCherry-P2A-LAT-EGFP were added to the imaging chambers with different pMHC densities and incubated for 30 min at 37 °C. Then the cells were fixed by adding the same volume of ice-cold 4% paraformaldehyde in PBS (2% final concentration) and incubating for 10 min at room temperature. The chambers were washed by PBS and imaged within 1 day. Cells with Cal520 were also treated and fixed with the same procedures, then the cells were permeabilized by 0.33% Triton-X 100 in PBS for 5 min and washed by PBS. Permeabilization washed out Cal520 to make LAT-EGFP-positive cells distinguishable. It also improved the preservation of cell morphology presumably due to the release of osmotic pressure. The cells without Cal520 were also permeabilized in some of the replicates, but it did not noticeably change the result. LAT-EGFP-positive cells with adhesion were selected and imaged with 488 nm excitation TIRF (100 ms, 2.6 $W/cm^2$) and 561 nm epi-fluorescence (100 ms, 3, 4, and 5 μm) channels with 1.5x magnification tube lens. Most cells exhibited either clearly cytosol-localized or nucleus-localized NFAT-mCherry, consistent with previous reports (*41*, *68*), which were manually classified. About 30-50 cells were observed for each chamber to measure the proportion of the activated cells (nucleus-localized) with 95% confidence interval (Clopper-Pearson). Rare population with unclear localization was omitted from the analysis. The obtained dose-dependence curves were fit by four-variable sigmoid curves (upper and lower bound, $EC_{50}$, Hill coefficient) with least squares regression.

For the cells with Zap70-mNeonGreen-P2A-LAT-mCherry and mNeonGreen-Vamp7-P2A-LAT-mCherry, the positive cells were found and imaged within 20 min after the cell addition, and the cells that are already adhering to the SLB were also included for imaging. The images were acquired with 640 nm excitation TIRF (500 ms, 3.3 W/cm$^2$), 561 nm excitation TIRF (200 ms, 6.7 W/cm$^2$), 488 nm excitation TIRF (100 ms, 2.6 or 4.9 W/cm$^2$ for Vamp7, 2.6 W/cm$^2$ for Zap70) channels with 1.5x magnification tube lens and the frame interval of 3 sec.

For the Cal520 bleaching measurement, the cells were imaged with 488 nm excitation epi-fluorescence (20 ms, 4 μm) channel with 1x magnification tube lens and the frame interval of ~20 ms (minimal dead time).

For the pMHC bleaching measurement, SLB with high-density pMHC (100% labeled MCC-Atto647N, 14 μm$^{-2}$) was imaged with 640 nm TIRF (500 ms, 3.3 W/cm$^2$) with 1x magnification tube lens with the frame interval of ~500 ms (minimal dead time).

The Calcein measurements were performed with the same conditions as the corresponding Cal520 measurements.

For the imaging of Fab'-ligand binding to Jurkat cells, the cells loaded with Cal520 and expressing LAT-mCherry were added to the bilayer functionalized with human ICAM-1 and Fab' ligands and imaged with 640 nm excitation TIRF (500 ms, 3.3 W/cm$^2$), 561 nm excitation TIRF (100 ms, 3.6 W/cm$^2$), and 488 nm excitation epi-fluorescence (20 ms, 4 μm) channels with 1.5x magnification tube lens. For the imaging of LAT condensates and calcium signal, the cells loaded with Cal520 and DiD and expressing LAT-mCherry were imaged with 640 nm excitation TIRF (50 ms, 3.3 W/cm$^2$), 561 nm excitation TIRF (100 ms, 3.6 W/cm$^2$), and 488 nm excitation epi-fluorescence (20 ms, 4 μm).

pMHC density determination

The density of pMHC molecules were determined by the single-molecule images with 640 nm TIRF channel with short exposure time (20 ms, 92 W/cm$^2$). The images were cropped to extract the central region with nearly homogeneous illumination intensity, then analyzed with Trackmate(*85*) to detect the spots (LoG detector, 0.2 μm radius, no median filter, with subpixel localization, manually optimized threshold). The density was determined from the number of detected spots and the total area analyzed. The density up to about 0.4 μm$^{-2}$ gave the isolatable single-molecule spots and could be determined by this method. The higher densities (up to ~5 μm$^{-2}$) were determined by the linear extrapolation of intensity-density relationship from lower densities. The partially labeled pMHC molecules were used for higher densities than ~5 μm$^{-2}$, for which the total pMHC density was calculated based on the labeled fraction. The density of Fab' ligands were also determined by the same methods.

Image processing and cell segmentation

The images with LAT, pMHC, Cal520 channels were background-subtracted and corrected for illumination intensity heterogeneity. The background signal was measured with a dark-control chamber which contains only the imaging buffer on glass coverslip or SLB. The illumination intensity map was measured with the solution of fluorescein, rhodamine B, or 3,3′-diethylthiadicarbocyanine iodide (DTDCI) for each wavelength (compounds from Sigma

Aldrich). The images with RICM channel were background-subtracted with manually optimized background level and each pixel intensity was converted to the absolute value for the following segmentation process. The stage movements during the image acquisitions to track the cells, if any, were corrected so that the image cross-correlation is minimized for the pre- and post-movements. The images with LAT, RICM, and Cal520 channels were then segmented for each cell by frame-by-frame thresholding as follows. The images were gaussian-blurred (gaussian kernel $\sigma$) for segmentation. The region fully inside the cell ($R_{in}$) and the region fully encompassing the cell ($R_{out}$) were manually selected for each frame, which were typically simple circular shapes. The mean intensity in $R_{in}$ was measured ($I_{in}$), and the segment within $R_{out}$ was obtained with threshold $\alpha_{low} I_{in} < I < \alpha_{high} I_{in}$ where $\alpha_{low}, \alpha_{high}$ is manual factors and $I$ is pixel intensity. The segment was refined with binary opening with the square kernel ($K_{opening}$), then the regions with the sizes smaller than a manual minimum ($A_{min}$) were removed. Multiple regions were allowed to exist after these processes, which was rare cases. The parameter values for the segmentation procedures are shown in **Table S1**. The example images for the segmentation procedures are shown in **Fig. S24**. The frame-by-frame threshold enabled consistent segmentation especially for Cal520 images which exhibit variable intensities.

The area of cell segments from LAT or RICM channels were used to define the cell adhesion timings. The adhesion event was defined as the timing when the area exceeds and remains above a threshold for a certain duration (20 μm$^2$ and 20 frames for LAT, 10 μm$^2$ and 10 frames for RICM). The time axes are normalized to the adhesion timing throughout the analyses unless otherwise stated.

<u>pMHC:TCR complex detection</u>

The pMHC:TCR complexes were detected using Trackmate (version 6.0.1) (*85*). The extended-exposure TIRF images after background subtraction, illumination correction, and image shifting correction were analyzed with Trackmate to detect the spots (LoG detector, 0.2 or 0.3 μm radius, no median filter, subpixel localization, manually optimized threshold). The detected spots inside the cell segment as well as the spots close enough to the segment (less than 11 pixels away) were selected. The remaining spots were tracked (simple LAP tracker, 1 μm max linking distance, no gap closing), and mislinkings and misdetections were manually corrected.

<u>LAT condensates detection</u>

LAT condensates were also detected with Trackmate. The images from LAT channels after background subtraction, illumination intensity correction, and image shifting correction were gaussian blurred (σ: 1 pixel) and the condensates were detected (LoG detector, 0.3 μm radius, no median filter, subpixel localization, manually optimized threshold). Unlike pMHC:TCR, the detected spots outside of the cell segment as well as the spots near the cell periphery (less than 11 pixels away) were deleted. This segment-filtering reduced the false-detection at cell periphery. The remaining spots were tracked (simple LAP tracker, 1 μm max linking distance, no gap closing), and mislinking and misdetection events were manually corrected. The condensates detected for only single frame were omitted from analyses to avoid the potential short condensate-like artifacts such as membrane fluctuations. For the co-imaging of LAT with Zap70 or Vamp7, the condensate tracks with three or more frames were considered, reflecting the faster frame rate for these measurements. For the ratiometric detection with Jurkat cells, LAT and DiD intensity images were normalized to the whole-time whole-cell average, gaussian-blurred (σ: 1 pixel), and converted to the ratiometric images. The region with normalized DiD signal above

0.1 was considered. The condensate detection was performed in the same way as LAT images for primary T cells.

Detection of pMHC:TCR-proximal LAT condensates

pMHC:TCR-proximal LAT condensates were detected based on the existence of colocalized pMHC:TCR at the moment of condensate formation essentially as described previously (*49*). If pMHC:TCR exists within 0.25 µm of the condensate at the first frame of the condensate track, it was considered pMHC:TCR-proximal. For the Zap70 and Vamp7 measurements, if pMHC:TCR exists within 0.4 µm at the first and second frames, the condensate was considered pMHC:TCR-proximal. Modified criteria were used for these measurements to take advantage of the faster frame rate.

Cal520 signal quantification

Normalized intensity change ($\Delta I/I_0$) was calculated as the following. After background subtraction and illumination intensity correction, the pixel intensities within the cell segment were converted to the 5th-95th percentile average. The lowest and highest 5% pixels were omitted for the robustness against potential artifacts from low- and high-intensity objects. The intensity was corrected for the experimental bleaching curve. The first 5 frames were averaged to yield $I_0$, and $\Delta I/I_0 = (I - I_0)/I_0$ was calculated.

Cal520 photo-bleaching correction

The empirical bleaching curve was obtained by imaging the cells with Cal520 on ICAM-only supported bilayer with fast frame rate (~20 ms interval). The intracellular region was manually selected for each cell and the mean intensity traces were obtained, then normalized to the initial intensities (first 5 frames). The population-averaged normalized intensity trace, representing the photo-bleaching decay, was obtained. For the bleaching correction of data, the measured Cal520 intensity trace (frame-wise) was divided by the empirical normalized bleaching curve. We note that the bleaching curve under $Ca^{2+}$ saturation condition was also measured for the cells with ionomycin and extra $CaCl_2$ added during the measurement (condition described above). The bleaching curve obtained under the $Ca^{2+}$ saturation condition exhibited minor differences from that of the basal condition (**Fig. S1**). The bleaching curve for the saturation condition was fit well by a single-exponential decay, while the curve for the basal condition was only fit well by double-exponential decay. We used the bleaching curve from the basal condition for data correction, since the LAT-$Ca^{2+}$ co-imaging data typically exhibited the $\Delta I/I_0$ of about 0-3, which is much closer to basal condition ($\Delta I/I_0$ of about 0) than saturation condition ($\Delta I/I_0$ of about 11).

Cal520 response calibration

The *in situ* dynamic range of the Cal520 was evaluated by measuring Cal520 intensities from the cells under $Ca^{2+}$ depletion (BAPTA-AM and calcium-free buffer), basal (ICAM-only bilayer), and $Ca^{2+}$ saturation conditions (ionomycin and extra $Ca^{2+}$). Basal and saturation conditions were measured in the same experiment by imaging the cells before and after the drug addition. The cells were added to the imaging chamber for each condition and the snapshots of multiple cells were taken within 10 min. The mean Cal520 intensity at depletion ($I_{depletion}$) and saturation ($I_{saturation}$) conditions were used to construct the response curve using the dissociation constant of Cal520 from the manufacturer ($K_d$ = 320 nM), assuming the following relationship:

$$I = I_{depletion} + \left(I_{saturation} - I_{depletion}\right) \times \frac{[Ca^{2+}]}{[Ca^{2+}] + K_d}$$

, where the depletion condition represents the non-specific signal without $Ca^{2+}$. The intensity from the basal condition was treated as $I_0$ to construct the relationship between $\Delta I/I_0$ and [$Ca^{2+}$].

Cross-correlations and response functions

The $N_{condensate}$ trace was defined as the momentary number of LAT condensates co-existing at each frame. The condensate size $I_{condensate}$ was defined for each condensate as follows:

$$I_{condensate}(t) = A_{in} \frac{I_{in}(t) - I_{out}(t)}{Cell\ Average}$$

where $I_{in}$ and $I_{out}$ are the mean intensities in the condensate circular region (0.5 μm radius) and the condensate periphery donut-shaped region (0.5-1.0 μm radius), $A_{in}$ is the area of the condensate circular region in the unit of pixels, and $Cell\ Average$ is the mean pixel intensity of the cell segment for all frames (**Fig. S21A**). $\Sigma I_{condensate}$ was defined as the sum of $I_{condensate}$ for the co-existing condensates for each frame. The traces were resampled with the interval of 0.1 s (nearest-interpolation for $N_{\text{condensate}}$, linear-interpolation for $\Sigma I_{condensate}$) before the cross-correlation calculation.

The cross-correlation $C_{xy}(\tau)$ of the traces $x(t)$ and $y(t)$ were defined as Pearson's correlation coefficient with lag:

$$l_\tau := \begin{cases} \{0, 0.1, \dots, t_{max} - \tau\}\ (\tau \geq 0) \\ \{-\tau, -\tau + 0.1, \dots, t_{max}\}(\tau < 0) \end{cases}$$

$$C_{xy}(\tau) = \frac{Mean[x(t)y(t+\tau)] - Mean[x(t)] \times Mean[y(t+\tau)]}{Std[x(t)] \times Std[y(t+\tau)]}\ for\ t \in l_\tau$$

where $t_{\text{max}}$ was set to be 600 s. The LAT traces were substituted for $x$ and $\Delta I/I_0$ was substituted for $y$, thus the peak at positive lag would indicate that the Cal520 trace is delayed from LAT traces. The false-correlation control was evaluated as the cross-correlation between the traces from different two cells (performed by cycling the cell indices). This false correlation will reflect the global patterns consistent among cells, if any. The resulting cross-correlation traces were evaluated as the mean and SD.

The response curves were constructed by plotting the $N_{condensate}$ - $\Delta I/I_0$ value pairs from each frame, pooled from all cells. To correct the delay between the image acquisitions for different channels, $\Delta I/I_0$ was linearly interpolated at the acquisition timings of LAT image. The frames after 0 s and before 600 s were used.

Onset analysis

The onsets for LAT condensates and calcium signal were determined by change point detection using a Python library ruptures (*86*). The momentary number of LAT condensates $N_{condensate}$ and $\Delta I/I_0$ were resampled with 0.1 sec interval (nearest-interpolation for $N_{condensate}$ and linear interpolation for $\Delta I/I_0$). The traces were zero-padded (100 seconds) at the beginning for the detection of the onset at early timepoints. The change points of the mean were determined by PELT algorithm with manually optimized penalty values for each variable (same values among cells, typically 0-6 change points per trace), and the first change points were defined as the onsets. Cal520 onset was not clearly seen in the cells with low signal, thus the cells with the total cumulative $\Delta I/I_0$ lower than 50 [sec] were omitted. For the cells with both onsets detected, correlation was analyzed by linear regression after omitting the detection outliers (two-tailed Grubbs test for the onset difference, α=0.05). The examples of the detected change points are shown in **Fig. S7**.

<u>Spectral analyses and impulse response estimation</u>

Impulse response was estimated under LTI system approximation. The system output $y(t)$ represents $\Delta I/I_0$ and the input $x(t)$ represents formation events, $N_{condensate}$, or $\Sigma I_{condensate}$, which are connected by the convolution of impulse response $h(t)$, and $y(t) = x(t) * h(t)$. The empirical transfer function was determined by:

$$H = \frac{\langle CPSD \rangle}{\langle PSD(x) \rangle} = \frac{\langle Y\bar{X} \rangle}{\langle X\bar{X} \rangle}$$

where the capital letters represent the Fourier transform, brackets represent population-average, and bars indicate complex conjugate. For this calculation, the data was resampled with the interval of 0.1 s. For the LAT condensate formation events, which is essentially a point process, the input was expressed as a set of gaussian spikes (σ = 1 s). The input and output within the time window [0, 600] sec was converted to the estimated CPSD and PSD by discrete Fourie transform with Hanning window function. To consider the Nyquist frequency of the time resolution of measurements (6-7 sec intervals), the spectra were lowpass-filtered (<0.07 Hz). The transfer function *H* and impulse response $h$ were calculated. The main peak in the impulse response was found near zero second. Impulse response was cropped with the window [-20, 60] sec to extract the main peak, and the corresponding transfer function was recalculated from the cropped impulse response.

To evaluate the expected stochastic fluctuations, the formation events following Poisson process was simulated, and the resulting output traces were calculated by the convolution of the cropped impulse response. The PSD of the simulated output traces were calculated in the same way as the experimental output traces. The experimental PSD was shown as logarithmic mean and standard deviation among population, and the simulated PSD was shown as the logarithmic mean among sufficiently large number of simulation trials (1000 trials). The rate of Poisson process for this simulation was determined as the population-average of the experimental formation rate, which was calculated by the whole-time (0-600 sec) formation frequency for each cell.

The rationale for using population-average rate is as follows. First, cell-to-cell variation of the underlying Poisson process rate $r_i$ ($i$ is cell index) apparently exists (cumulative formation events are apparently more spread than stochastic noise) but it is only less than a factor of about 2 (**Fig. 7A, S20G**). Thus, only a first-order correction would be necessary. Second, the output PSD of LTI-system for the input following a Poisson process with rate *r* follows $\langle PSD(y) \rangle = r|H|^2$ for the asymptote of long observation time (Poisson noise or shot noise). Thus, the expected stochastic fluctuations expressed as PSD are approximately proportional to the rate for all frequencies. From those,

$$\frac{\langle PSD(y_i) \rangle_i}{\langle simulated\ PSD(r_i) \rangle_i} \approx \frac{\langle PSD(y_i) \rangle_i}{simulated\ PSD(\langle r_i \rangle_i)}$$

represent the deviation of experimental PSD from simulated Poisson noise, where $\langle \cdot \rangle_i$ denotes the population average. Therefore, it is reasonable to use the population-average of the experimental formation frequency as the best estimate of $\langle r_i \rangle_i$. We used the numerically simulated PSD instead of the analytical approximation $r|H|^2$ for better accuracy, though the difference was confirmed to be small (**Fig. S25**).

<u>Detection of Zap70 and Vamp7 colocalization with LAT condensates</u>

For Zap70 measurement, the cells transduced with Zap70-mNeonGreen and LAT-mCherry co-expression construct with P2A sequence were applied onto SLB with pMHC (MCC-Atto647N) molecules and imaged with TIRF for Zap70, LAT, and pMHC channels. Due to the low transduction efficiency, the positive cells were searched for after applying the cells, and the cells that are already adhering to the SLB were also included for imaging. The imaging was performed within about 20 min after cell addition to minimize the cells undergoing signaling events for too long duration. Zap70 images exhibited small membrane-bound clusters and the strong and smooth background of cytosolic Zap70 signals. Due to the low intensity with fluctuations, individual Zap70 clusters could not be tracked. Alternatively, LAT condensates were tracked first, then the existence of colocalized Zap70 cluster was determined for each condensate track. The condensates with the track length longer than 3 were considered in this analysis. After the background correction, illumination correction, and gaussian-blur ($\sigma$=1 pixel), cytosolic background was removed by white tophat filter with 5-pixel square kernel (**Fig S26A**). The Zap70 cluster was detected based on local signal-to-noise frame-by-frame as follows. The local background signal $I_{BG}$ was calculated as the average pixel intensity in the donut-shaped periphery region around the condensate location. The periphery region was defined by the radius larger than the periphery inner radius (0.4 µm) and smaller than the periphery outer radius (1.0 µm) (**Fig S26B**). The local background noise $\sigma_{BG}$ was calculated as the robust estimate of standard deviation ($0.7413 \times (75\%\ Quantile - 25\%\ Quantile)$) in the periphery region. Within the detection radius (0.25 µm) around the condensate location, if any of the pixel intensity exceeded $I_{BG} + (3.5 \times \sigma_{BG})$, the Zap70 cluster was considered detected for that frame. Then, if the colocalized Zap70 cluster was detected for more than two consecutive frames, the corresponding condensate was considered as Zap70-positive. These detection criteria were determined so that the resulting classification agrees with visual inspection.

For Vamp7 measurement, the cells transduced with mNeonGreen-Vamp7 and LAT-mCherry co-expression construct with P2A were imaged in the same way. Vamp7-associated vesicles exhibited transient signals in the TIRF imaging plane with low background signals. The colocalized Vamp7-vesicles were detected for each condensate in the same way as Zap70 with a few modifications: the white tophat filtering was not performed for Vamp7, the periphery inner radius was changed to 0.5 µm, and the intensity threshold was changed to $I_{BG} + (3.0 \times \sigma_{BG})$.

### Supplementary Text

Supplementary Text S1. Observations of Jurkat cell behaviors.

We set up the imaging system for Jurkat cells, a commonly used model cell line, using the bilayer-tethered Fab' ligand system which we previously developed and characterized for murine T cells (*61*, *81*). Fab' ligands were prepared from anti-human CD3 IgG (OKT3 clone). Jurkat cells transduced with LAT-mCherry adhered well to the bilayer functionalized with human ICAM-1. Fab' ligands (0.01-0.1 $\mu m^{-2}$) bound strongly to the TCR on the cells (**Fig. S2A**). Most cells exhibited cell membrane ruffling, which obscured the detection of LAT condensates. To overcome this difficulty, the cells were pre-treated with membrane-stain DiD as a counter stain and three-color imaging was performed for LAT-mCherry, DiD, and Cal520 channels (**Fig. S2B**). Fluorescence signal from Fab' ligands were negligibly small compared to bulk DiD signal and were not observed for this measurement. The ratiometric image of LAT and cell membrane stain was used for the detection of LAT condensates. The obtained traces of the momentary number of LAT condensates ($N_{\text{condensate}}$) and the calcium signal did not exhibit noticeable correlations (**Fig. S2C**). There were rare population of cells (about 5 to 20% by visual inspection with the microscope) which exhibited smooth cell membrane with no ruffling, and these cells exhibited apparently correlated fluctuations (**Fig. S2D, E**). However, we could not implement the experiments to analyze these rare cells with sufficient throughput in an unbiased way. The membrane ruffling and uncorrelated fluctuations indicate the altered signaling behaviors in Jurkat cells. Similar membrane ruffling has been reported, and it has also been reported that Jurkat exhibits distinct cytoskeletal behaviors (*87*, *88*). Hyperactivity of calcium signaling-related molecules have also been reported (*89*, *90*). Due to these behaviors, which were not observed with primary murine T cells, Jurkat cells appeared unsuitable model for our purpose.

Supplementary Text S2. Extended description of onset analysis.

As described in the main text, we observed the simultaneous onsets of LAT condensation and calcium signaling. We also observed, although qualitatively, the morphological kinapse-synapse transition occurring simultaneously. Mechanistic knowledge implies the mutual dependences among three events: LAT condensate induces $Ca^{2+}$ elevation, kinapse-synapse transition depends on $Ca^{2+}$ and LAT-mediated signals (*64*), and kinapse-synapse transition potentially affects TCR triggering by transforming the membrane tension and the distribution of signaling molecules (*65*, *66*). Potentiation of T cell signaling by kinapse-synapse transition might occur through such concerted phenomena.

The onset timings were quantified by change-point detection. In most cases (6/10 cells for lower density and 5/6 cells for higher density), both $N_{\text{condensate}}$ onset and Cal520 onset were clearly discernable and could be detected (**Fig. S7A**). In some cases (4/10 cells for lower density and 1/6 cell for higher density), calcium signal is too low overall and the calcium onset was not discernable (**Fig. S7B**). These cells were omitted from the onset analysis by applying a certain threshold. In rare cases (1/10 cell for lower density), the onset was barely discernable by visual inspection and the change-point detection was not accurate (**Fig. S7C**). These cases were filtered by statistical outlier detection (Grubbs test) applied to the difference of onset timings (**Fig. 5**).

We note that the positive cross-correlation between $N_{\text{condensate}}$ and calcium signal traces were not a mere reflection of the simultaneous abrupt rises at the onsets. It was shown by calculating cross-correlation of the traces shifted by the onset (**Fig. S8**). These shifted traces have the calcium onset at time zero (the traces without calcium onset are not shifted). The false-correlation control, the cross correlation between different cells, was still near zero and the true cross-correlation was significantly higher. This suggests that the cross-correlation dominantly reflects the fluctuations following the onsets and not the abrupt rises at the onsets themselves.

The observation of condensation and calcium onsets well-separated from cell adhesion (delayed by 0 to 150 sec) was unique to this experiment with low-density pMHC. Several studies have reported the delayed calcium rise using high-dose stimulatory surface, using wide-field fluorescence microscopy which is not capable of distinguishing the cell contact and adhesion and spreading. In these cases, the measured delays represent the contact-onset delay, which ranges from 30 sec to 10 min with the typical values of several minutes (*5*, *9*, *11*, *18*, *91–93*). The studies which specifically measured adhesion-onset delays, typically using TIRF microscopy, reported shorter delays ranging from 7 to 30 sec (*46*, *94*, *95*). The studies which used APCs to stimulate the T cells and observed contact-onset delays also reported long delays ranging from 3 min to 10 min (*16*, *96*, *97*). A study which used APCs to stimulate the T cells and observed the delays using differential interference contrast microscopy, which allows the observation of cell adhesion to some degree, reported shorter delays ranging from 40 to 150 sec (*9*). A study which used photo-activatable pMHC that allows the initiation of pMHC:TCR ligation after the completion of adhesion reported much shorter adhesion-onset delay, 6.5 sec (*94*). These previous studies indicate long delays between cell contact and cell adhesion, which is inferred to be several minutes. We observed a delay between cell-SLB contact and adhesion in some cases as mentioned in the main text, typically ranging from 0 to 5 minutes, which is consistent with the inferred value. Kinapse-synapse transition also occurs immediately and simultaneously with calcium elevation upon contact with high-dose stimulatory surface (*63*). The present work and those reports combined suggest that the adhesion-onset delay is only several seconds with high-dose stimulations, but it gets extended to several minutes with low-dose stimulations.

<u>Supplementary Text S3. Extended description of the results for anti-TCR-coated glass condition</u>

High-density anti-TCR-coated glass coverslips were prepared by incubating 10 μg/mL of anti-mouse TCR IgG (clone H57), following previously reported methods (*70*). The primary AND-TCR T cells transduced with LAT-mCherry and loaded with Cal520 were added upon the coverslip and imaged. LAT immediately formed condensate throughout the interface, which dissolved in about 1 min, and the $Ca^{2+}$ signal showed an immediate rise and the following slow decay over more than 10 min (**Fig. S10A, B**). For low-density anti-TCR-coated glass coverslips, 1000-fold lower concentration of anti-TCR at 10 ng/mL was used for incubation. With the low-density condition, a wide range of responses were observed: some cells exhibited non-isolatable LAT condensation throughout the interface and $Ca^{2+}$ rise and decay, similar to high-density anti-TCR. Some cells exhibited weak overall response characterized by poor adhesion, few or no LAT condensates, and weak or no $Ca^{2+}$ elevations. Most notably, the rest of the cells exhibited the modest overall response characterized by the formation and dissolution of isolated LAT condensates, whose formation events persisted for more than 10 min (**Fig. S10C-E**). The $Ca^{2+}$ signal in these cells exhibited the temporal patterns with an initial rise and following fluctuations, which resembles the cells with pMHC stimulations. The diverse response to low-density anti-TCR is possibly due to the heterogeneous density of the antibody on the glass substrate which may be difficult to control.

<u>Supplementary Text S4. Uncertainty evaluation of the number of detected pMHC:TCR-proximal LAT condensates</u>

The number of detected pMHC:TCR-proximal LAT condensates (9 ± 6%, mean ± SD) would underestimate the actual number if fluorescently labeled pMHC is photo-bleached significantly. The measured photo-bleaching time-constant for pMHC (labeled with Atto647N) was 589 frames (**Fig. S11C**). LAT-mCherry and Cal520 co-imaging data typically contained about 100 frames until 10 min after the cell adhesion. Therefore, the population of photo-bleached pMHC molecules will reach up to about 16%. Since the detected pMHC:TCR-proximal condensates were 9% on average, the photo-bleached pMHC:TCR-proximal condensates are estimated to be $0.09 \times \frac{0.16}{0.84} = 1.7\%$ as the upper bound. The Zap70 and Vamp7 measurements typically contained about 200 frames for 10 min acquisition, with which the photo-bleached pMHC molecules will reach up to 29%. Given that the 4 - 10% of the condensates were detected as pMHC:TCR-proximal, the photo-bleached pMHC:TCR-proximal condensates were estimated to be 1.6 - 4% as the upper bound. In all these cases, the photo-bleached pMHC:TCR-proximal condensates are much less than the distal condensates, which account for about 90%.

The detection sensitivity of pMHC:TCR fluorescence signal also affects the detection of proximal condensates. In the procedures of pMHC:TCR detection using TrackMate (see Materials and Methods), detection threshold values were manually optimized. After this automatic spot detection, further manual correction was performed. The manual correction was very little and the number of detected proximal condensates was unaffected (**Fig. S12**). For the detection threshold values, if the threshold is systematically low or high and false-positive and false-negative detections occur, the detection of proximal condensates is affected. We varied the threshold values and tested the detection of proximal condensates (**Fig. S12**). With much lower threshold (-100), pMHC:TCR detection was very sensitive and a lot of false-positive detections occurred. The number of detected pMHC:TCR increased to about 278% of the original condition, yet the number of detected proximal condensates increased to only about 135%. With much higher threshold (+100), some pMHC:TCR were clearly missed. The number of detected pMHC:TCR decreased to about 71%, while the number of detected proximal condensates decreased to about 77%. In both cases, the number of detected proximal condensates was only slightly susceptible to the detection accuracy of pMHC:TCR spots, with the relative uncertainty of less than about 20%.

### Supplementary Text S5. Extended description of the results for Zap70 and Vamp7 imaging

To observe the presence of Zap70 in LAT condensates, Zap70-mNeonGreen and LAT-mCherry were co-expressed in T cells and imaged with TIRF on SLB with labeled pMHC (**Fig. S13A**). Zap70 exhibited small transient clusters with fluctuating intensities that occasionally colocalized with either pMHC:TCR-proximal or distal LAT condensates. Due to the small cluster size, the tracking of Zap70 clusters was not feasible. Thus, we performed the Zap70 cluster detection at each LAT condensate trajectory based on the local intensity enrichment (see Materials and Methods). Zap70 clusters were detected in both pMHC:TCR-proximal and distal condensates with modest fractions (**Fig. S13B, C**).

To observe the recruitment of LAT-containing vesicles with Vamp7, mNeonGreen-Vamp7 and LAT-mCherry were co-expressed and imaged along with pMHC:TCR. The blobs of Vamp7 transiently coming to the TIRF illumination field were observed, indicating the Vamp7-containing vesicles recruited to the plasma membrane (**Fig. S14A**). Vamp7 signal preferentially appeared at the center of the cell-SLB interface, consistent with the enrichment near MTOC as previously reported (**Fig. S15**) (*75*). Due to the difficulty in tracking the transient blobs, the colocalization was analyzed for each LAT condensate trajectory in the same way as Zap70.

Colocalization of LAT and Vamp7 was partial: some of the Vamp7-containing vesicles and some of the LAT condensates colocalized (**Fig. S14B, C**), which is also consistent with previous studies (*75*). pMHC:TCR-distal condensates with colocalized Vamp7-containing vesicles accounted for 19% of the total condensates (**Fig. S14C**). Those condensates possibly include the plasma-membrane LAT condensates further recruiting vesicular LAT (*78*) or only the vesicular LAT without condensates, which were indistinguishable in our imaging resolution. pMHC:TCR-proximal condensates with colocalized Vamp7 were rare, which may include coincidental colocalizations (**Fig. S14B, C**).

Supplementary Text S6. Extended description of the spectral analyses.

The stochastic properties of condensation phenomena were analyzed to probe the stochastic effects on the fluctuation behaviors. For lower pMHC density ($0.10 \pm 0.03\ \mu m^{-1}$), once they begin, LAT condensation events occurred at apparently constant frequencies with whole-time-average frequency (0-600 sec) of $0.07 \pm 0.03\ s^{-1}$ (mean ± SD) (**Fig. 7A**). The intervals among formation events were approximately exponentially distributed, and the sequence of intervals exhibited no observable autocorrelations, indicating the independence among intervals (**Fig. S19**). These characteristics suggest that condensate nucleation approximates a stationary Poisson process. Condensate lifetime and size exhibited no population-wide time-dependence during the observation for both densities (**Fig. S20E, F**). Lifetime and size were also independently distributed among condensation events with no observable autocorrelation (**Fig. S19**). These properties suggest that the formation timing, lifetime, and size of the condensates are randomly distributed with little temporal heterogeneities, and their intrinsic stochastic variations drive a large part of the fluctuations in the momentary number of condensates ($N_{condensate}$) or the summed size of condensates ($\Sigma I_{condensate}$, **Fig. S21**).

From the dataset with higher pMHC density ($1.1 \pm 0.2\ \mu m^{-2}$), the condensate formation events exhibited non-uniform rates over time in some cases, with the whole-time-average frequency of $0.08 \pm 0.03\ s^{-1}$ (mean ± SD) (**Fig. S20G**). On population-average, the formation events were most frequent at about 200 sec after adhesion, then decreased to a similar rate with low pMHC density (**Fig. S20D**). $N_{condensate}$, $\Sigma I_{condensate}$, and calcium levels exhibited a peak at the same timing, reflecting this frequent formation events **(Fig. S20A-C**). This time-dependence was not so strong that the interval statistics differs from stationary Poisson process, as the interval distribution was also approximately exponentially distributed and positive autocorrelation was not observed (**Fig. S19**). These behaviors indicate that slow variations in the condensation frequency start to appear as the pMHC density increases over this density. But at this density, the stochastic fluctuations are still relatively large, and the formation events are not exactly the same as but approximating a Poisson process in terms of the degree of stochastic fluctuations.

We analyzed the relation between LAT condensates and calcium signal by LTI-system approximation. A minimal model considering only the formation timings of condensates is represented by the LTI-system with the point-process input of formation events and the output of calcium signal, so that each formation event triggers a fixed pulse (impulse response) of calcium signal. Two other models considering condensate lifetime (input of $N_{condensate}$) and condensate size (input of $\Sigma I_{condensate}$) were compared. $N_{condensate}$ model represents the situation where each condensate is continuingly inducing calcium signals over its lifetime. $\Sigma I_{condensate}$ model represents the situation where each condensate is continuingly inducing calcium signals proportional to its size. Data from all cells were pooled to determine the empirical impulse response for each pMHC density range. The impulse responses for $N_{condensate}$ or $\Sigma I_{condensate}$ models were short and immediate, while the impulse response from the formation events was more delayed and prolonged (**Fig. S17A-C**). Fine oscillations appeared in the impulse response functions due to limited time resolution of the measurements, but the different peak shapes were discernable. These differences successfully reflected the differences of three models: tight correlation between $N_{\mathrm{condensate}}$ or $\Sigma I_{\mathrm{condensate}}$ and calcium signal suggests a delta-function-like impulse response, while the formation-events model is expected to exhibit the impulse response spanning over the pulse length of calcium signal. The impulse responses independently obtained from the datasets with higher pMHC densities resembled those from lower pMHC densities, which cross-validates the reasonable estimation of impulse responses (**Fig. S18B**). As self-consistency tests, the convolution traces of the impulse response and the experimental input were compared to the experimental calcium traces. Three models recapitulated the experimental output similarly well, as the cross-correlations of experimental and convoluted traces were similarly positive (**Fig. 7C, S17D, S18A, C**). It was also consistently observed that using $\Sigma I_{\mathrm{condensate}}$ instead of $N_{\mathrm{condensate}}$ does not noticeably change the cross-correlation and the response function with calcium response (**Fig. S21, S4C-F**). Therefore, the condensate lifetime and size have minor influence on the overall calcium signal fluctuations, and formation-events model ignoring these factors still recapitulates the apparent behaviors of calcium signal fluctuations.

As a phenotypic description of calcium signal fluctuations, formation-events model with fixed pulses of calcium signals suffices. This model entails Poisson noise representing a purely stochastic component. To evaluate the contribution of stochasticity, the experimental power spectral density (PSD) of calcium traces and the expected PSD for Poisson noise were compared. Poisson noise was evaluated by simulating the formation events following a Poisson process and convoluting the empirical impulse response (see Materials and Methods). The simulated PSD matched the experimental PSD over a frequency range of about 5 to 20 mHz within a factor of about 5 (**Fig. 7D**). The same result was consistently observed for higher pMHC densities (**Fig. S18D**). Therefore, a significant portion of the calcium signal fluctuations can be described as originating from the stochastic variations of the condensate formation timings, without any underlying oscillatory biochemical circuits involved. It should be noted that this does not exclude the other two models as mechanistic possibilities. It is possible that condensates actively induce the calcium response over their lifetime with or without size-dependence, but the lifetime and size heterogeneity are only minorly influential on the overall fluctuation behaviors. In line with this scenario, cross-correlation analysis suggested that $N_{\mathrm{condensate}}$ was majorly determined by condensation timings rather than lifetime distributions (**Fig. S22**).

The time-dependent formation rate with a peak at about 200 sec observed with higher density is presumably reflected at lower frequencies than 5 mHz, although such a coherent and slow (compared to the observation time) component cannot be simply interpreted in this spectral analysis. This peak behavior might possibly encode the information of antigen density that determines NFAT translocation, since it was the only noticeable and deterministic difference between two tested densities. The titration experiment also exhibited the appearance of the calcium signal peak as the pMHC density increases over about 1 $\mu m^{-2}$, which supports this speculation (**Fig. 6**). We also note that we qualitatively observed a slow desensitization of calcium response per LAT condensate over time, which might possibly be some induced negative regulations (**Fig. S23**). This might cause some broadening in the frequency domain. The fluctuations with higher frequencies than 20 mHz which exceeds Poisson noise possibly include the measurement noise and the heterogeneity of calcium signal pulses which are averaged out in this analysis. The calcium traces are best described as these factors adding on top of the basal LTI-system-like stochastic fluctuations.

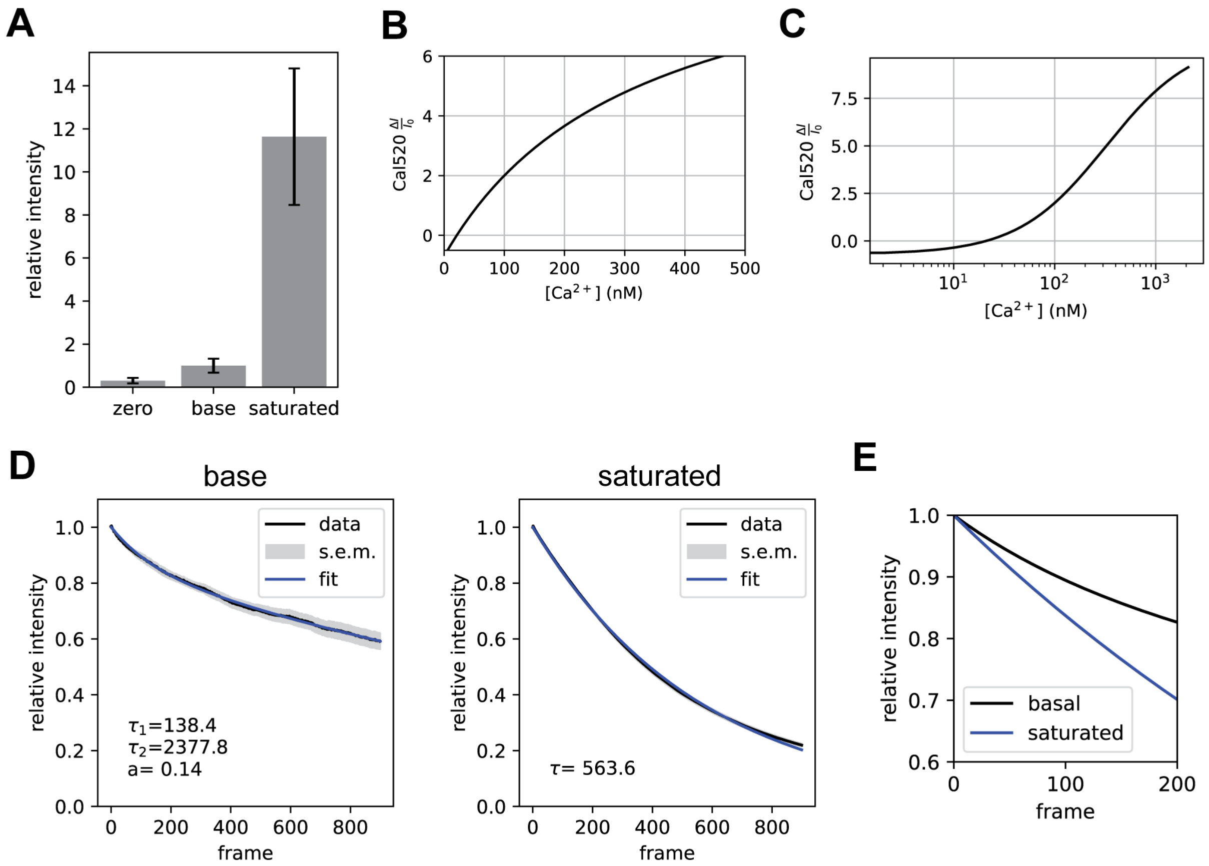


**Fig. S1. Validation of the calcium measurement by Cal520**

(**A**) The Cal520 intensity in the cells treated with calcium depletion (zero), basal (base), or calcium saturation (saturated) conditions normalized to the basal condition. Calcium depletion condition was achieved with BAPTA-AM preloading and calcium-free imaging buffer, and calcium saturation was achieved with high-concentration ionomycin and calcium ion. (35 cells for depletion, 8 cells for basal and saturation, error bars: SD) (**B, C**) The Cal520 response curve based on the dynamic range determined in (**A**), shown in normal scale (**B**) and log scale (**C**). (**D**) Experimental bleaching curve of Cal520 in the cells treated with basal or calcium saturation conditions (18 cells for basal and 22 cells for saturation conditions). Cal520 intensity was measured with fast frame rate (~20 ms per frame) to minimize the intracellular calcium concentration fluctuations, which was further canceled out by population averaging. Basal condition was fit by sum of two exponential decay $aexp\left(-\frac{t}{\tau_1}\right) + (1-a)\exp\left(-\frac{t}{\tau_2}\right)$, while saturated condition was fit by a single exponential decay $\exp\left(-\frac{t}{\tau}\right)$. (**e**) Comparison of two bleaching curves.

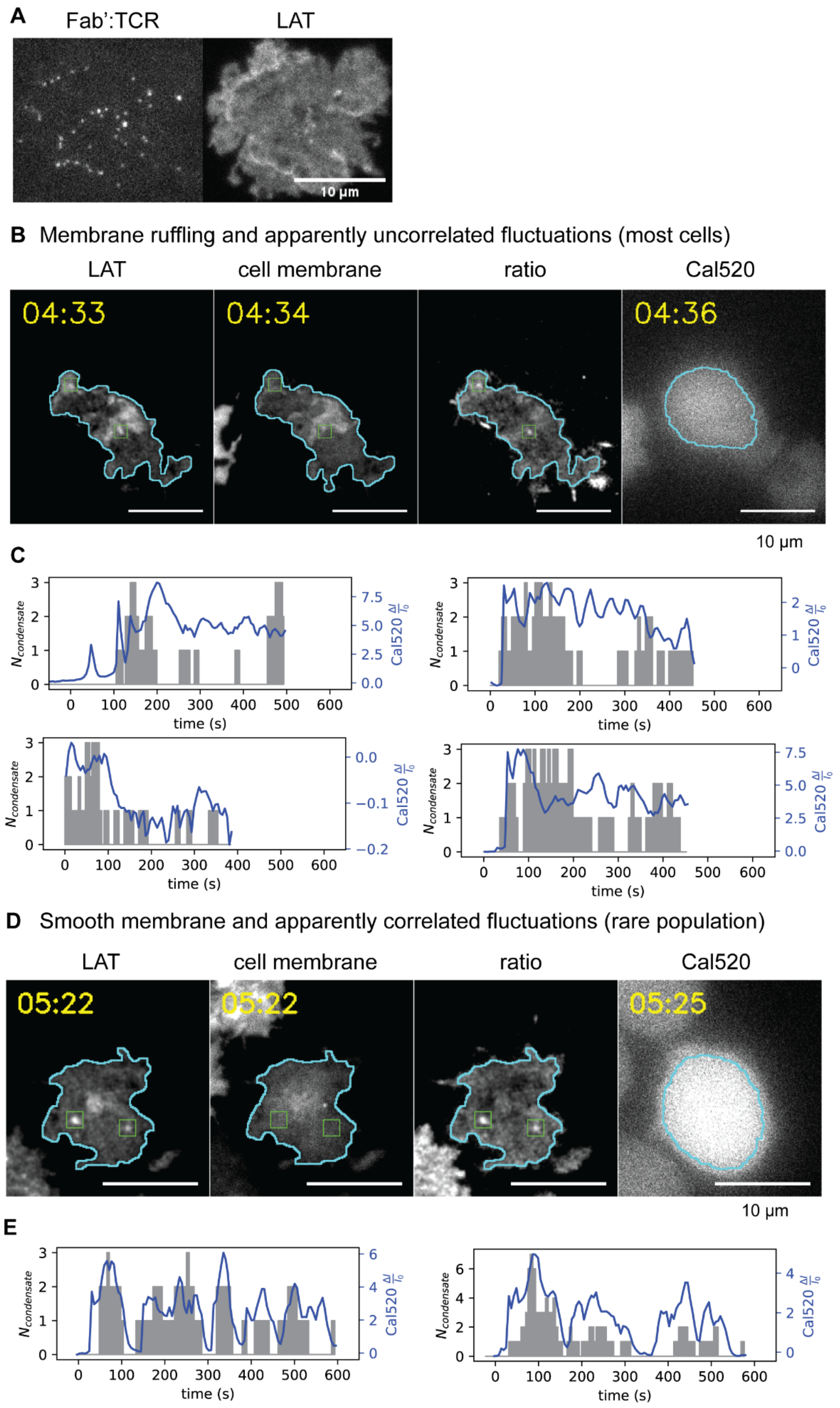
A
Fab':TCR
LAT
10 µm
B Membrane ruffling and apparently uncorrelated fluctuations (most cells)
LAT
cell membrane
ratio
Cal520
04:33
04:34
04:36
10 µm
C
$N_{condensate}$
Cal520 $\frac{\Delta I}{I_0}$
time (s)
D Smooth membrane and apparently correlated fluctuations (rare population)
LAT
cell membrane
ratio
Cal520
05:22
05:22
05:25
10 µm
E
$N_{condensate}$
Cal520 $\frac{\Delta I}{I_0}$
time (s)

**Fig. S2. Jurkat cell exhibited altered responses.**

(**A**) Anti-CD3 Fab' ligand (OKT3) tethered on the lipid bilayer (density of 0.09 $\mu m^{-2}$) were visualized with extended-exposure TIRF imaging. The Fab':TCR complexes underneath the Jurkat cell were selectively visualized as bright spots (left panel). Simultaneous TIRF imaging of LAT-mCherry visualized the cell adhesion (right panel). (**B-E**) Jurkat cells lenti-virally transduced with LAT-mCherry was stained with DiD (cell membrane stain) and loaded with Cal520, then imaged on Fab'-functionalized bilayer (density of 0.02–0.08 $\mu m^{-2}$). (**B, C**) Most of the cells exhibited the cell membrane ruffling and the ratiometric image was used to cancel out the ruffling and detect the condensates. Representative cell image is shown in **B**. In those cells, the traces of $N_{\text{condensate}}$ and calcium signal did not noticeably exhibit the correlated fluctuations. Observed traces for those cells are shown in **C**. (**D, E**) Rare population of the cells exhibited the smooth cell membrane and apparently correlated fluctuations.

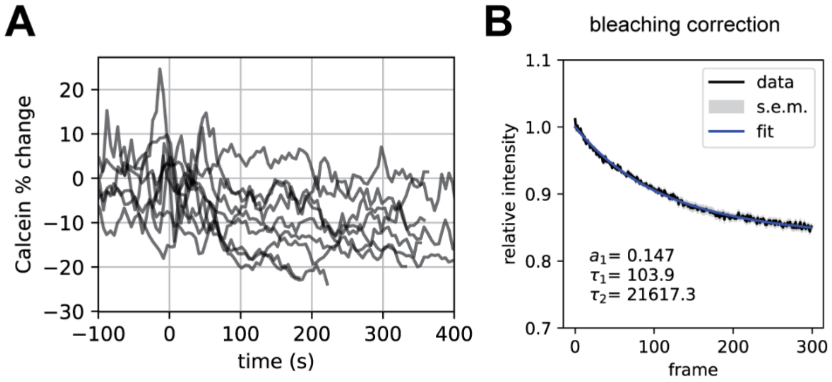


**Fig. S3. Evaluation of the cell-thickness-dependent intensity fluctuations by $Ca^{2+}$ non-responsive stain**

(**A**) Fluorescence intensity trace from the cells stained with Calcein. Intensity trace was normalized to the initial intensity, corrected for the bleaching curve, and shown as the percent change. The change was within about 20%. Data is pooled from 9 cells. (**B**) The experimental bleaching curve fit by $aexp\left(-\frac{t}{\tau_1}\right) + (1-a)\exp\left(-\frac{t}{\tau_2}\right)$, used for the bleaching correction. Bleaching curve is mean and s.e.m. from 24 cells.

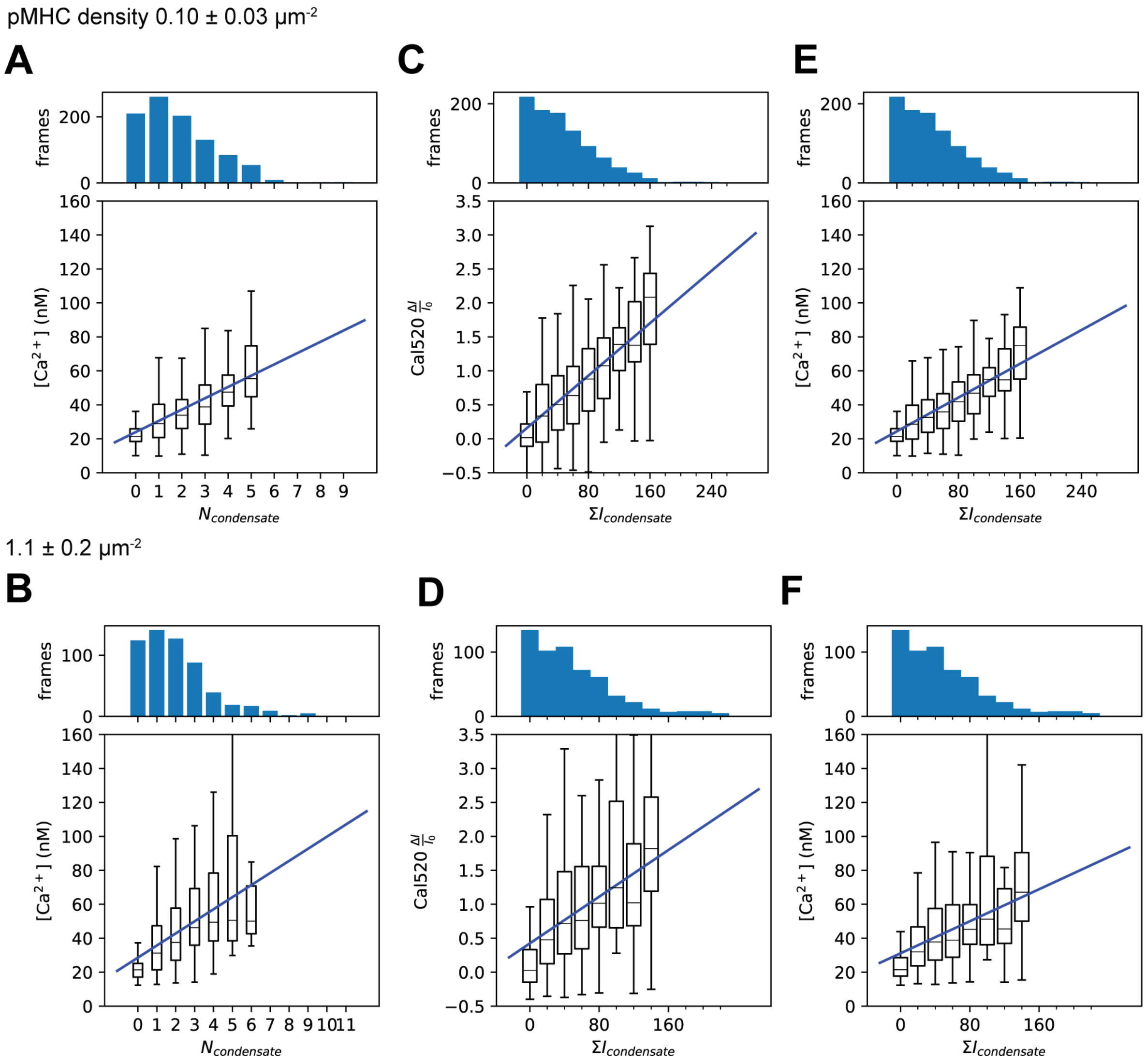


**Fig. S4. Response functions with different readouts**

Condensate-$Ca^{2+}$ response functions generated by framewise pooling for each pMHC density range. The readout of $Ca^{2+}$ signal was modified to the estimated absolute $Ca^{2+}$ concentration (**A**, **B**, **E**, **F**). The readout of the amount of condensates were modified to the summed condensate size (**C**-**F**).

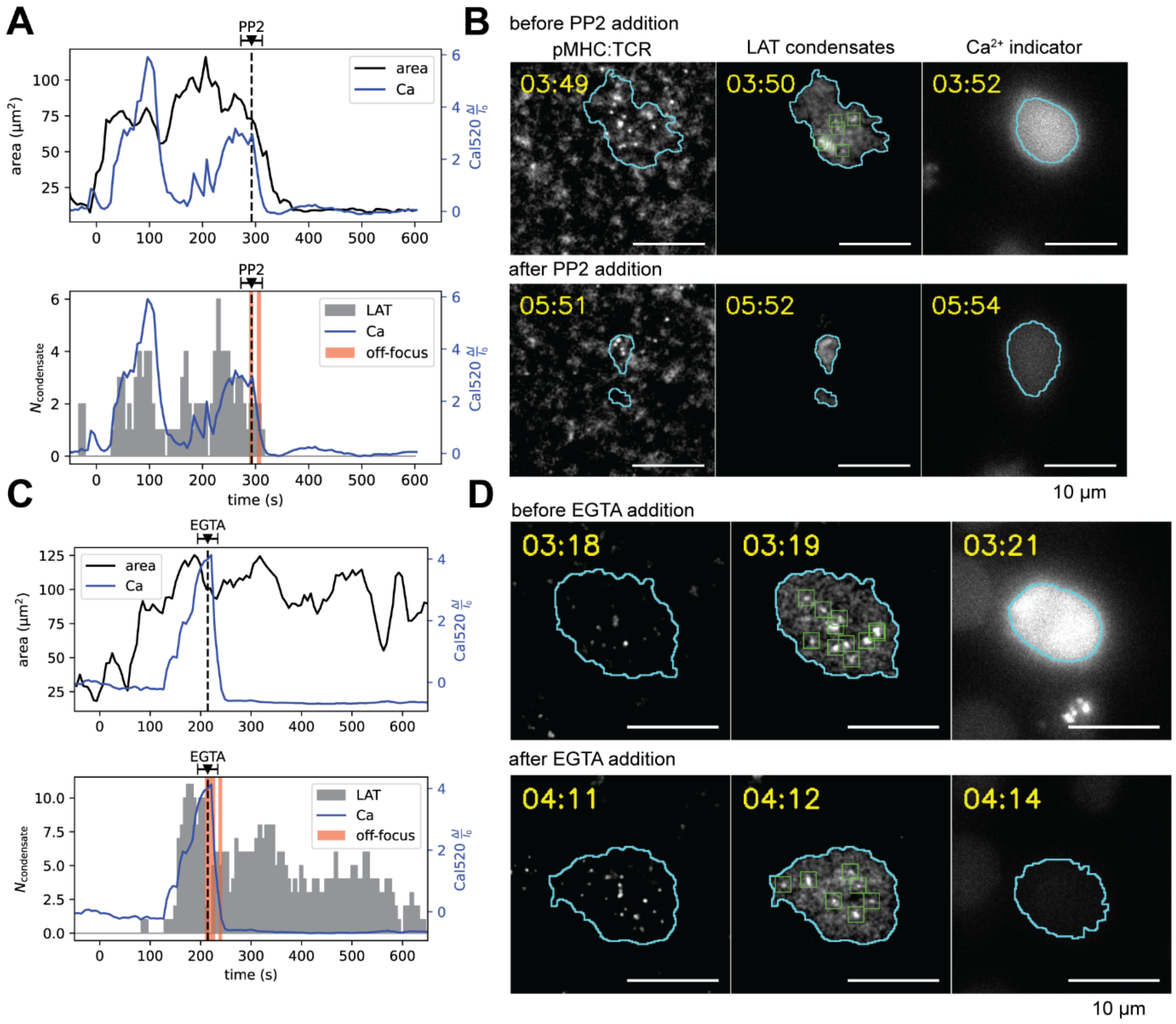


**Fig. S5. Validation of the direction of signal propagation with inhibitions**

(**A, B**) An example cell of LAT condensation inhibition experiment. T cell with LAT-mCherry and Cal520 was observed on SLB with 0.39-2.6 $\mu m^{-2}$ pMHC, and PP2 was added during the observation (timing annotated in **A**). LAT condensation and calcium signal were immediately quenched following the addition. While adhesion was also inhibited by PP2, pMHC still kept binding to the cell. Representative cell image from 3 cells, 2 mice is shown. (**C, D**) An example cell of extracellular calcium depletion experiment. EGTA was added during the imaging of the T cell on SLB with 0.24-1.8 $\mu m^{-2}$ pMHC to deplete the extracellular free calcium ion. The calcium signal immediately quenched after the addition, while LAT condensates kept forming. Some cells detached from the bilayer 1 to 5 min after EGTA addition, but the duration before the detachment was sufficient for the observation in most cases (5 out of 6 cells attempted). The data is representative from 5 cells, 3 mice. The manual addition of PP2 or EGTA solution involved the timing uncertainty of about ± 20 sec shown as error bars in **A**, **C**. A few frames of LAT images were not focused well to detect condensates due to drug addition procedures (red shades in **A, C**), but it did not obscure the overall observations.

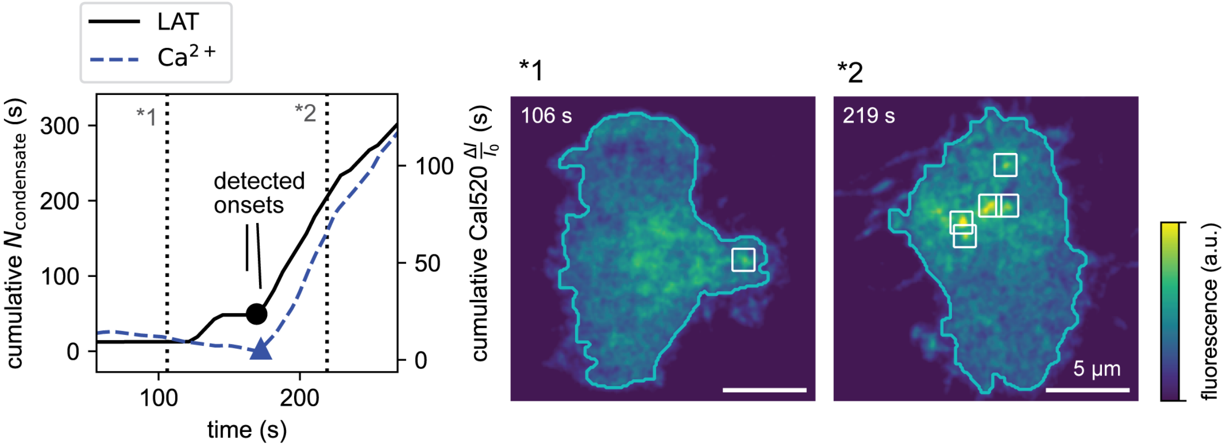


**Fig. S6. Additional example cell exhibiting simultaneous onsets**

(**A**) Cumulative traces of LAT condensate formation events and calcium signal. (**B**) Representative LAT snapshots before (*1 in **A**) and after (*2 in **A**) the onsets.

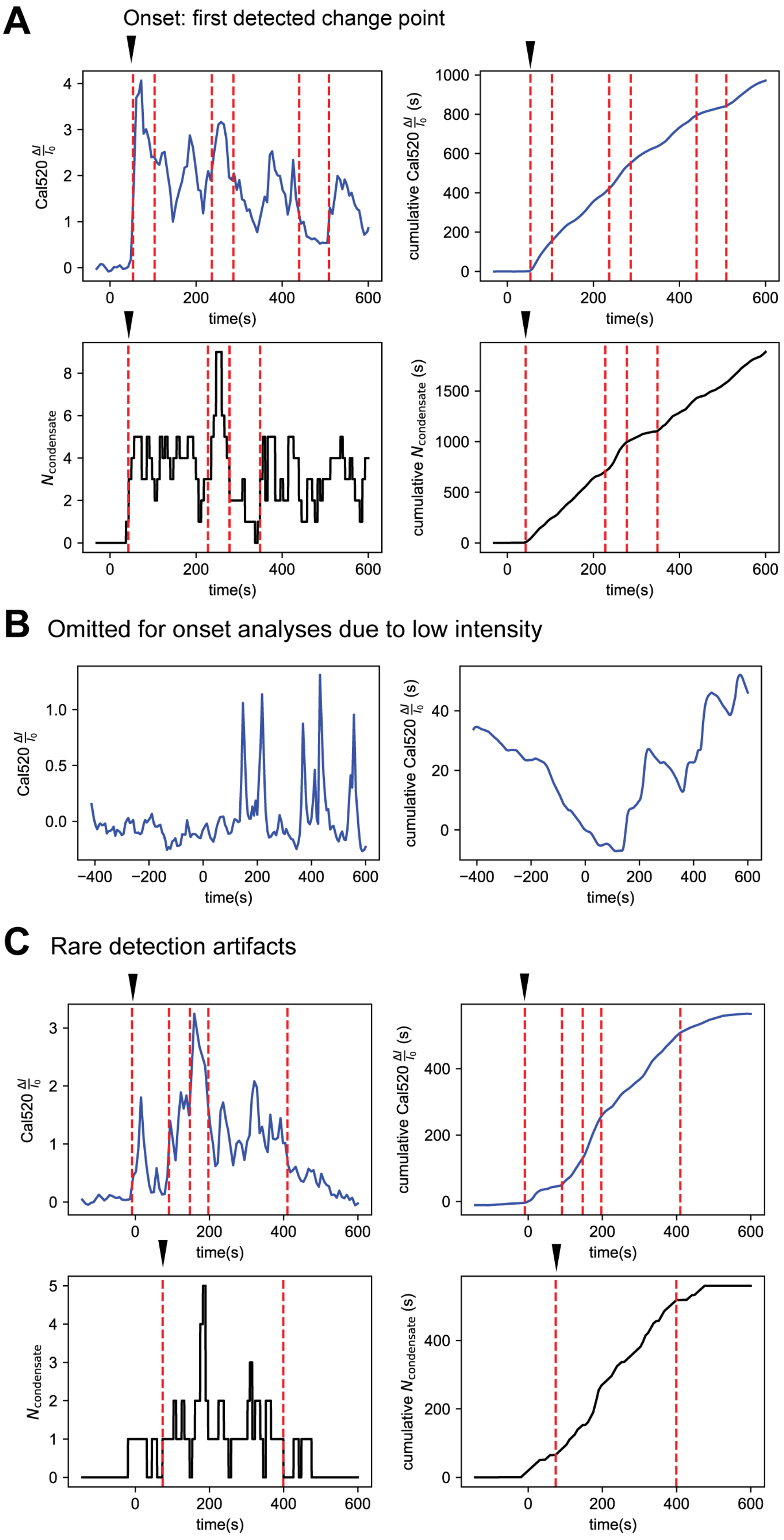
A
Onset: first detected change point
Cal520 $\frac{\Delta I}{I_0}$
time(s)
cumulative Cal520 $\frac{\Delta I}{I_0}$ (s)
$N_{condensate}$
cumulative $N_{condensate}$ (s)
B
Omitted for onset analyses due to low intensity
C
Rare detection artifacts

**Fig. S7. Quantification of the onset timings by change-point detection.** The onset timings for calcium levels and the momentary number of LAT condensates were detected by change-point detection. From the multiple change points detected (red dashed lines), the first change point was considered as the onset (black arrows). In most cases, the detected onset timings matched the visually discernable onsets seen in cumulative traces. A representative cell is shown in **A**. In some cells, the calcium levels were too low to discern the onset, as shown in **B**. (**C**) In rare cases, the onsets were detected in both LAT condensate and calcium signal by change-point detection, but the data were not sufficiently clear to define the onsets, resulting in detection artifacts.

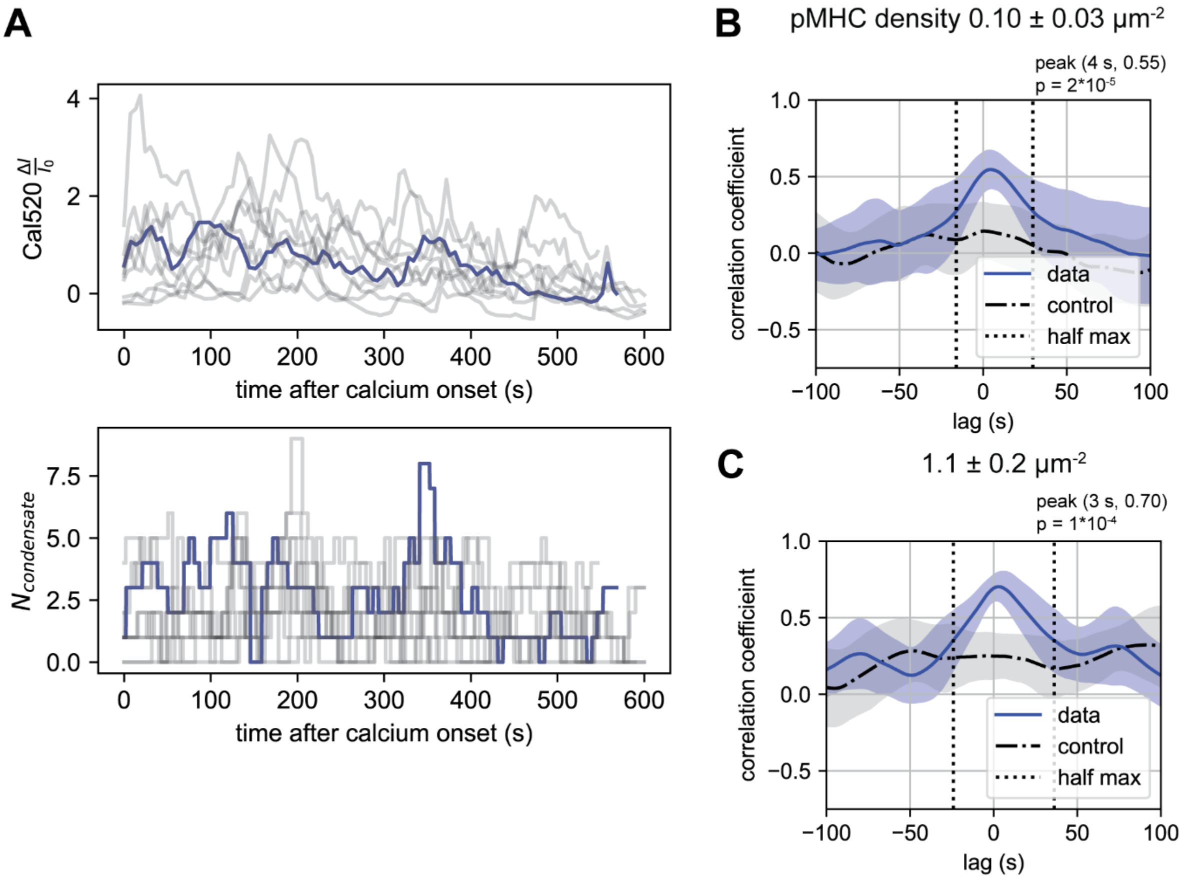


**Fig. S8. Simultaneous onsets have minor influence on LAT-calcium cross-correlation.**

(**A**) The example traces of Cal520 readout and $N_{\text{condensate}}$ with the time axis shifted to the calcium onset (0.10 ± 0.03 μm$^{-2}$ pMHC). The cells without detectable $Ca^{2+}$ onset were used without time shifting. Blue lines highlight an example cell. Fluctuations with cell-to-cell variations were observed after the onset. (**B, C**) Cross-correlation of time-shifted Cal520 readout and $N_{\text{condensate}}$ traces within 0-600 sec after the $Ca^{2+}$ onset. Significant cross-correlation was observed, showing the dominant effects from the fluctuations after the onsets. 7/10 cells for lower pMHC density and 5/6 cells for higher pMHC density exhibited detectable onsets of calcium signal.

**A**

| | $EC_{50}$ (µm$^{-2}$) |
|---|---|
| no treatment (n=3) | 0.6 ± 0.2 |
| with Cal520 (n=2) | 0.79 ± 0.09 |

**B**

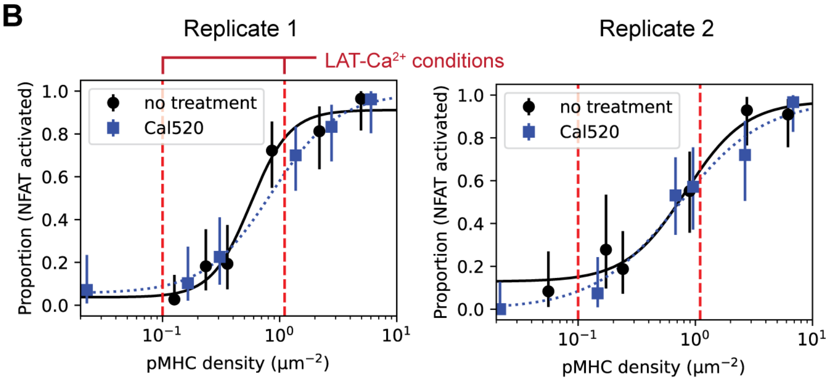


**Fig. S9. NFAT translocation assay**

(**A**) $EC_{50}$ pMHC densities (mean ± SD) for the conditions with and without Cal520 loading to the cells. n: the number of biologically independent replicates. (**B**) Additional data of the titration curves, showing two conditions overlaid for each experimental replicate.

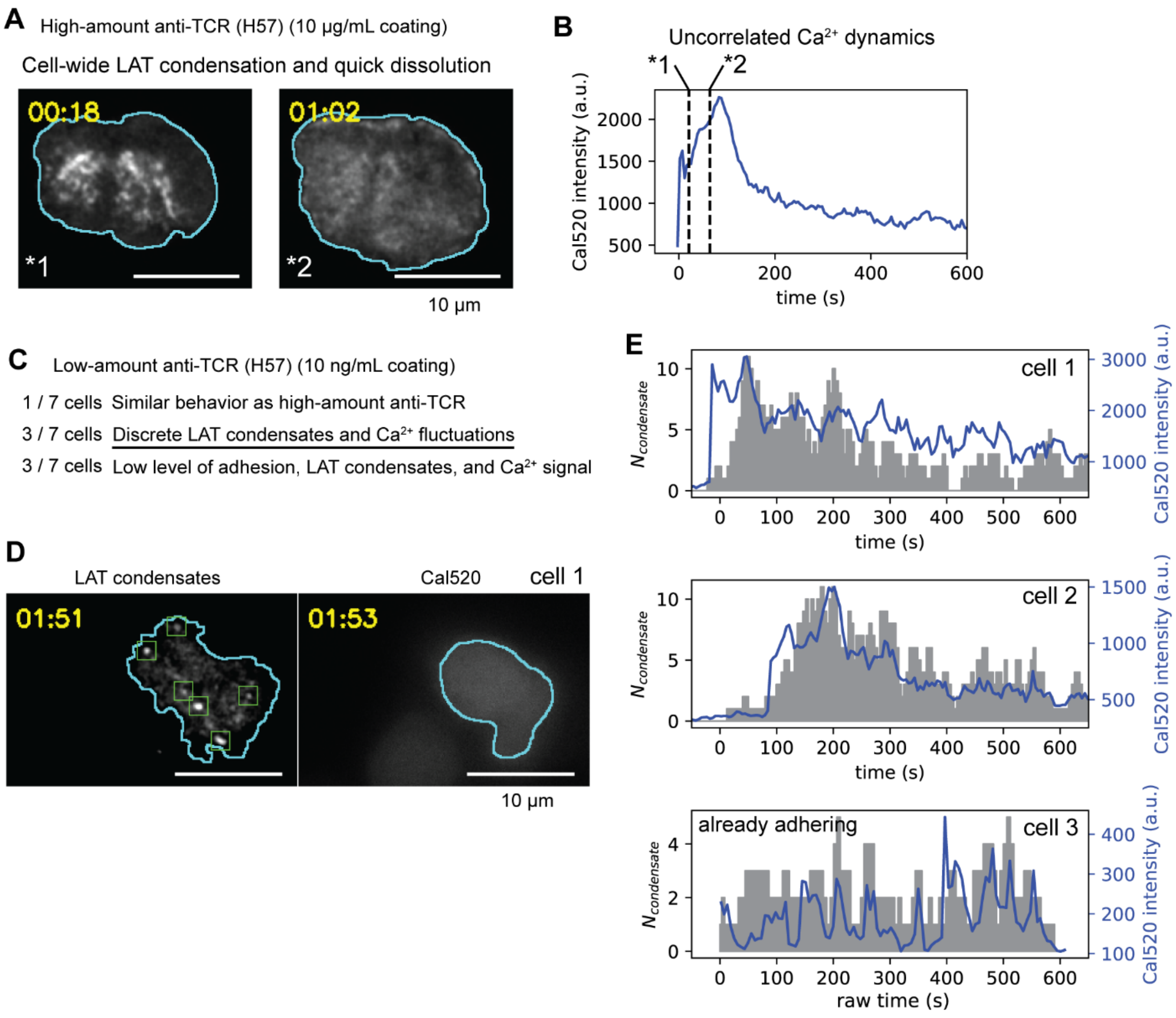


**Fig. S10. Surface-immobilized H57 anti-CD3 gave the temporal correlation.**

(**A, B**) T cells with LAT-mCherry and Cal520 were stimulated by high-density anti-TCRβ (H57) coated coverslips (10 μg/mL incubation). A representative cell from 3 cells, 2 mice is shown. (**A**) The corresponding LAT images showing the quick formation of superimposed LAT condensate (left) followed by quick dissolution (right). (**B**) Cal520 signal showed immediate rise-decay. Raw Cal520 intensity is shown due to too quick rise to determine initial intensity. *1 and *2 indicate the timings of the images in (**A**). (**C**) Low-density anti-TCRβ (H57) coated coverslips (10 ng/mL incubation) condition resulted in several distinct cellular behaviors. Some cells showed strong response similar to high-density condition, some cells showed modest response, and some cells showed weak adhesion, few or no LAT condensates and weak calcium response. 7 cells from 2 mice were imaged. (**D**, **E**) For the cells with modest responses, LAT condensates became discrete and kept forming for more than 10 min, allowing the quantification of $N_{\mathrm{condensate}}$ (**D**). The $N_{\mathrm{condensate}}$ and Cal520 signal exhibited fluctuating behaviors (**E**). The raw Cal520 signal traces are shown due to the difficulty in measuring the initial baseline intensity for a sufficient duration.

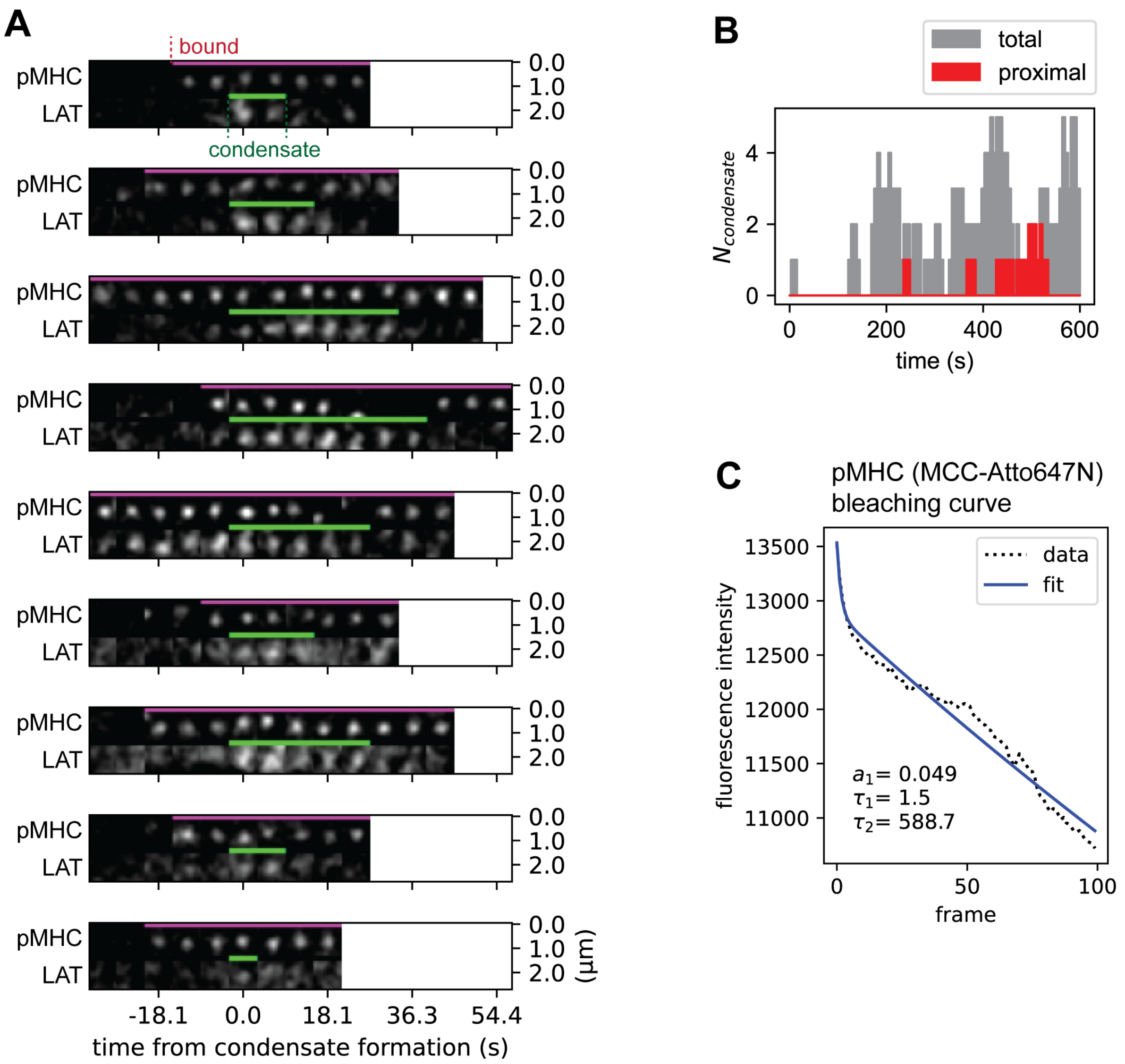


**Fig. S11. pMHC-TCR proximal LAT condensates were few.**

(**A**) Montages of the pMHC-TCR proximal LAT condensates from a representative cell. Images were centered at the location of the condensate or of the bound pMHC molecule when the condensate is absent. (**B**) The number of condensates trace from a representative cell. The traces accounting for the pMHC-TCR proximal condensates and the total condensates are shown. (**C**) The bleaching curve of pMHC (Atto647N-labeled MCC peptide, average of 4 measurements) was fit by sum of two exponential decays, $a_1 exp\left(-\frac{t}{\tau_1}\right) + (1 - a_1)\exp\left(-\frac{t}{\tau_2}\right)$. A small fraction of fast-bleaching component was presumably due to fluorescent contaminants. The bleaching time constant of pMHC was thus estimated to be $\tau_2 = 588.7$ frames.

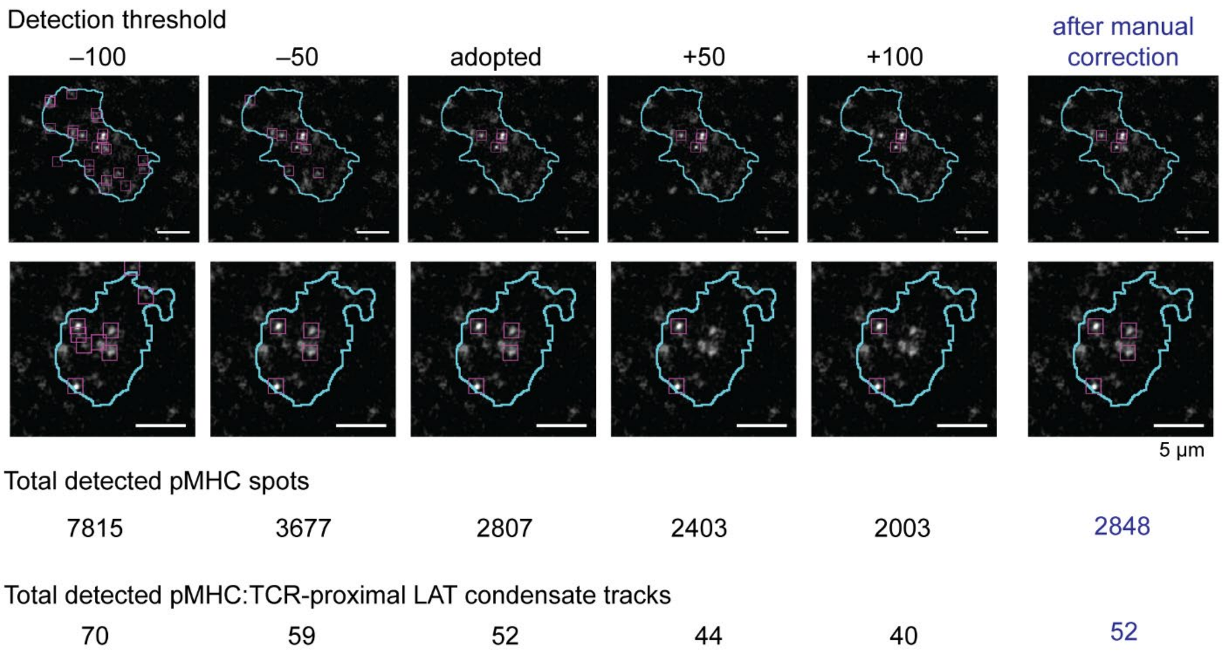


**Fig. S12. Detection of pMHC:TCR and proximal condensates with different threshold values.**

For the dataset with lower pMHC densities (0.10 ± 0.03 $\mu m^{-2}$), pMHC:TCR was reanalyzed with different detection threshold values in TrackMate. "adopted" indicates the threshold used in this study, and ±50 or 100 indicates the offset from that. In this study, the detected pMHC:TCR spots shown in "adopted" column were further manually corrected, which is shown in the "after manual correction" column. Top images are representative snapshots, and the detected spots are indicated as magenta boxes. Cell footprints are shown as cyan line. Note that the detection is limited underneath the cell (see Materials and Methods). Below images, the total number of detected pMHC spots (frame by frame) are shown. At the bottom, the total number of detected pMHC:TCR-proximal LAT condensate tracks are shown. "spot" refers to the detected object for each frame, and "track" refers to each molecule/object existing over multiple frames.

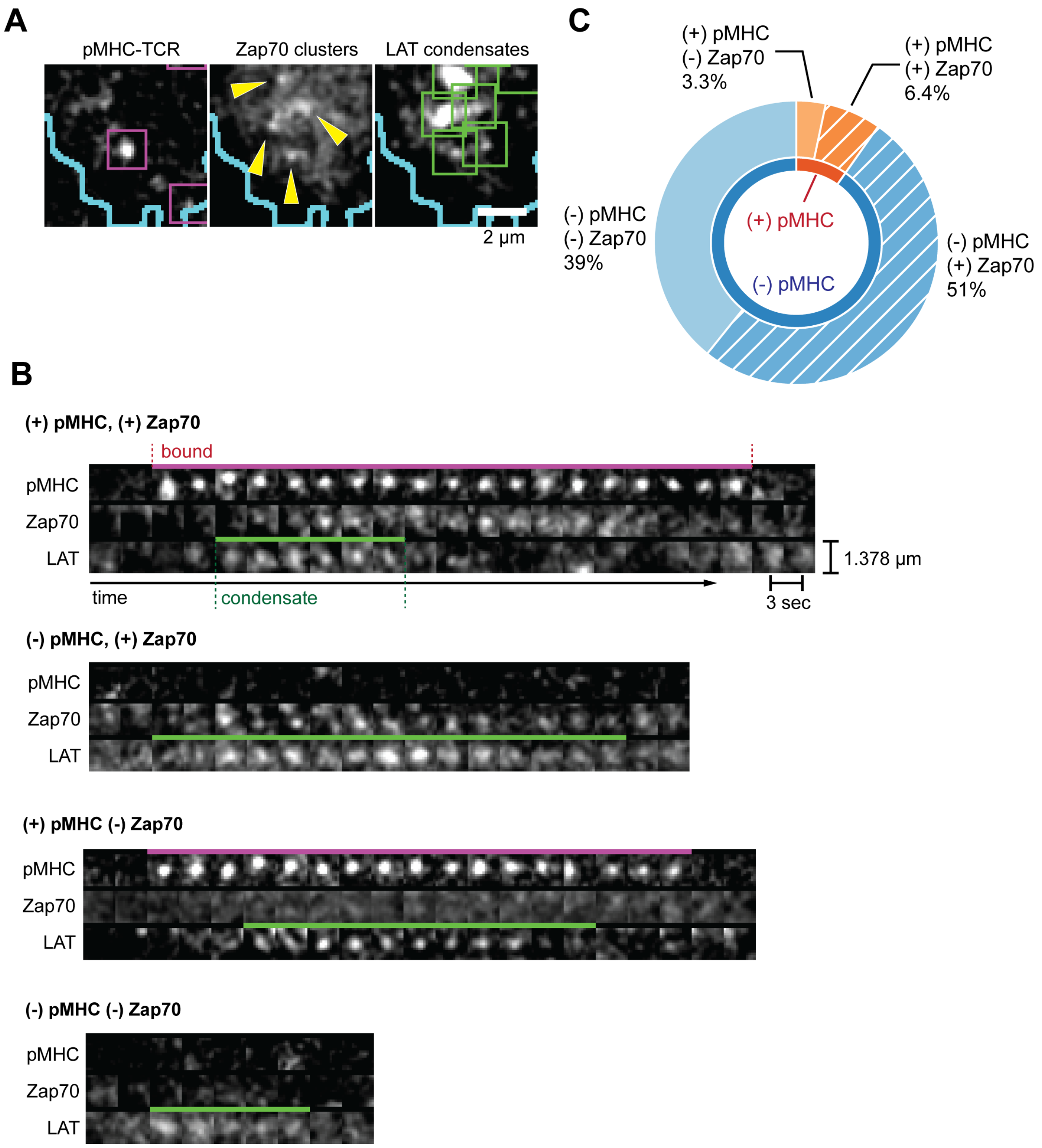


**Fig. S13. Detection of LAT condensates with colocalized Zap70 clusters**

(**A-C**) Colocalization of Zap70 clusters with LAT condensates were evaluated with living T cells expressing LAT-mCherry and Zap70-mNeonGreen on SLB with pMHC (0.18-0.25 $\mu m^{-2}$). LAT condensates were categorized based on whether the condensate formed at proximity of a pMHC-TCR complex and whether a colocalized Zap70 was observed. (**A**) An example cell snapshot showing visible Zap70 clusters (indicated by white arrows), some of which colocalized with LAT condensates. (**B**) Example montages of the four categories of LAT condensates. (**C**) The fractions of the condensates in each category (pooled from 6 cells from 2 mice, 451 total condensates).

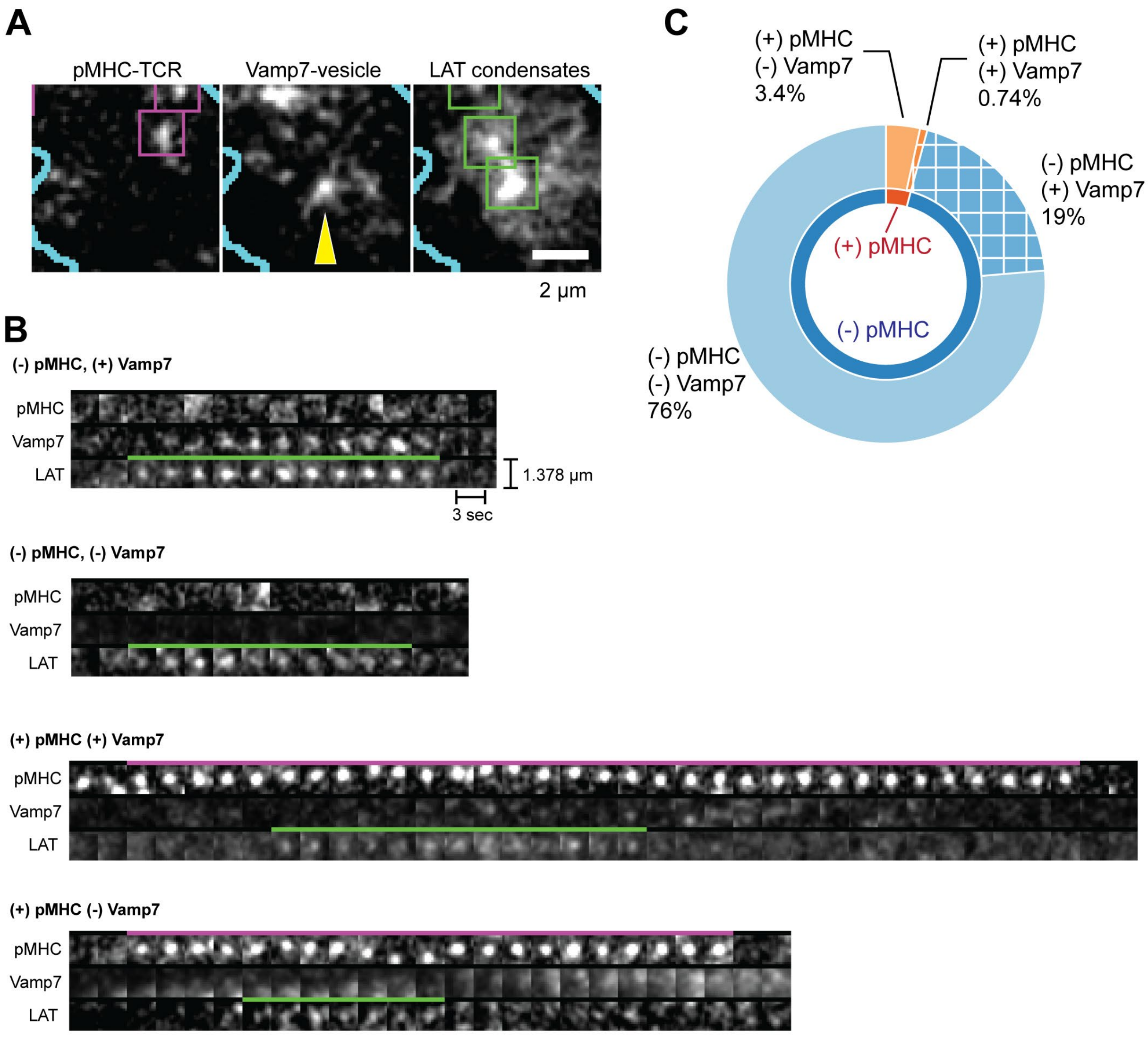


**Fig. S14. Detection of LAT condensates with colocalized Vamp7-associated vesicles**

(**A-C**) Colocalization of Vamp7-associated vesicles with LAT condensates were evaluated with living T cells expressing LAT-mCherry and mNeonGreen-Vamp7 on SLB with pMHC (0.19-0.29 $\mu m^{-2}$). LAT condensates were categorized based on whether the condensate formed at proximity of a pMHC-TCR complex and whether a colocalized Vamp7-associated vesicles was observed. (**A**) An example cell snapshot showing visible Vamp7-associated vesicles (indicated by white arrows), some of which colocalized with LAT condensates. (**B**) Example montages of the four categories of LAT condensates. (**C**) The fractions of the condensates in each category (pooled from 6 cells from 2 mice, 406 total condensates).

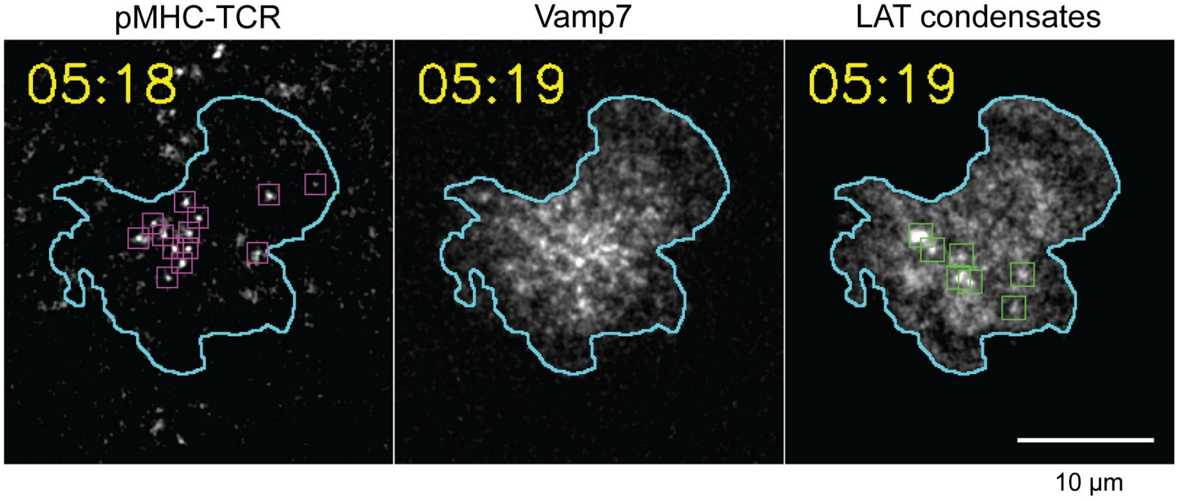


**Fig. S15. Representative cell snapshot of Vamp7 localization imaged with pMHC-TCR and LAT condensates.**

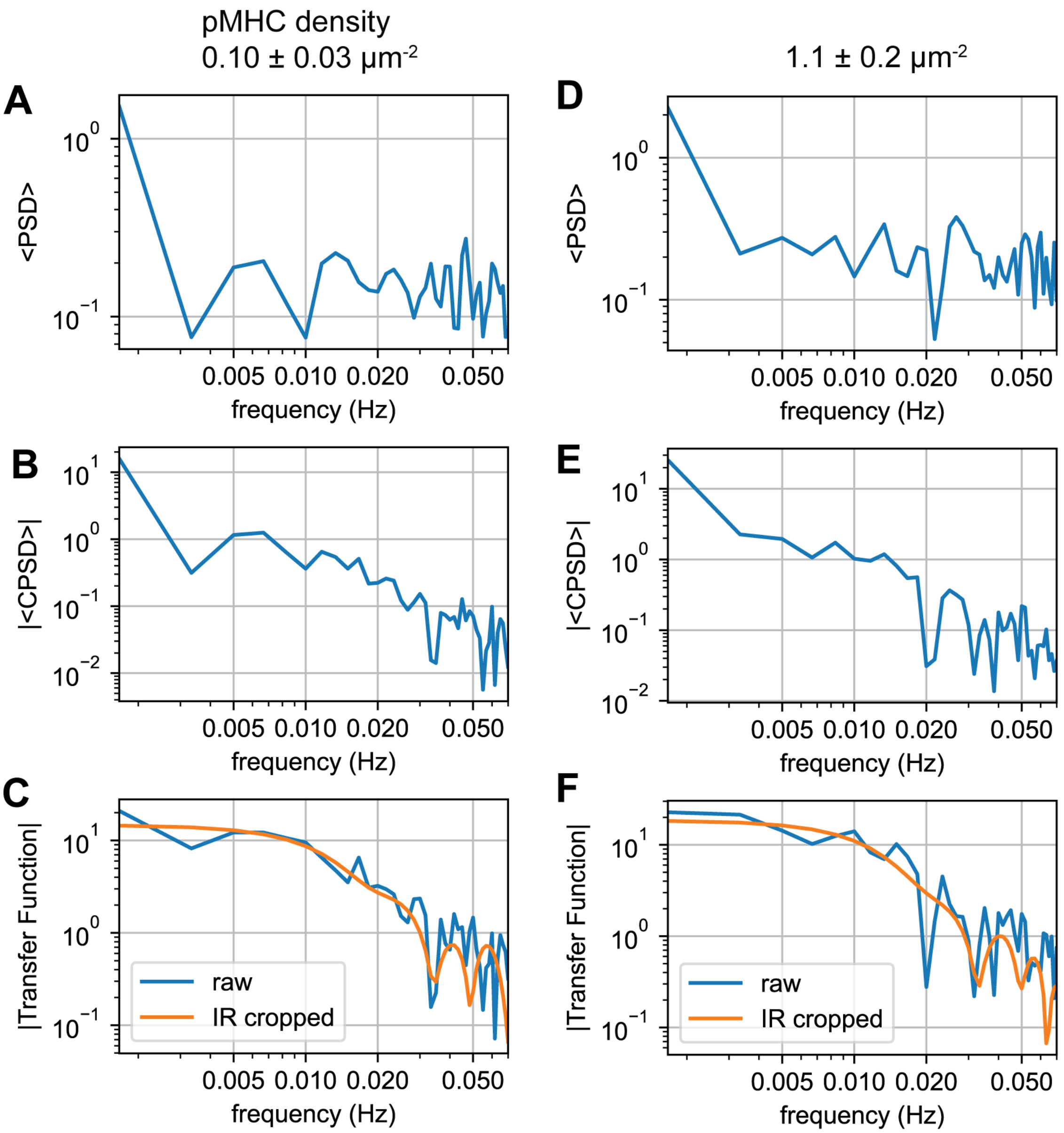


**Fig. S16. Spectral analyses for impulse response estimation for condensate formation events**

(**A, D**) Population-averaged power spectral density of the condensate formation events represented as a set of gaussian peaks. (**B, E**) Population-averaged cross-power spectral density of the condensate formation events and calcium signal. (**C, F**) Transfer function and the transfer function after cropping the impulse response. The dataset with lower pMHC density (0.10 ± 0.03 $\mu m^{-2}$) are shown in **A**-**C**, and the dataset with higher pMHC density (1.1 ± 0.2 $\mu m^{-2}$) are shown in **D**-**F**.

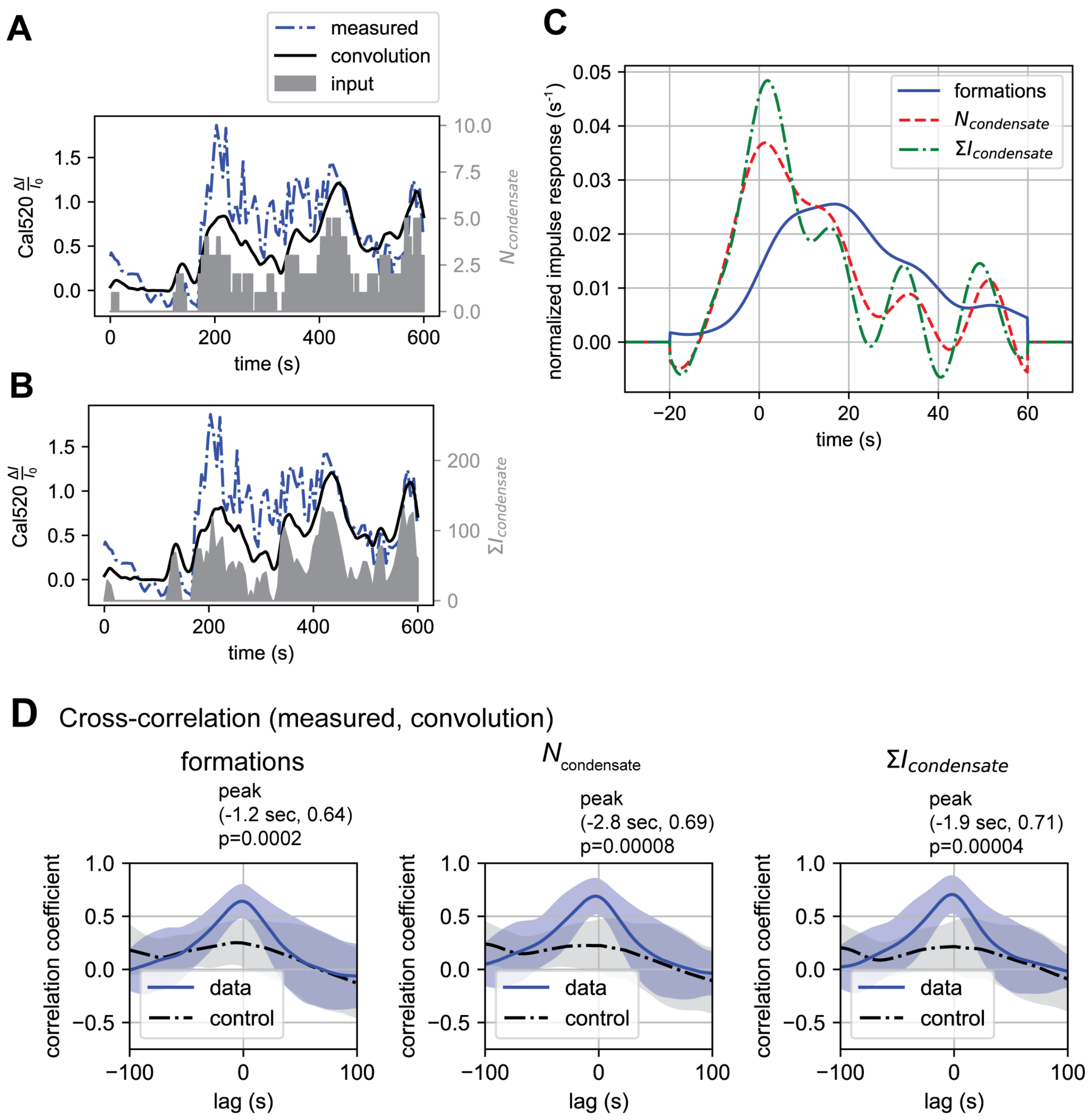


**Fig. S17. Impulse-response estimation for the continuous traces of LAT condensates**

(**A, B, C**) Impulse-response estimation using $N_{\text{condensate}}$ or $\Sigma I_{condensate}$ as the input instead of condensate formation events. The impulse-response for formation events showed delayed and extended response compared to the other two continuous inputs showing sharp and immediate peaks. Fine oscillation appeared due to limited time-resolution. (**C**). (**D**) Cross-correlation of the experimental Cal520 $\Delta I/I_0$ trace and the convoluted trace.

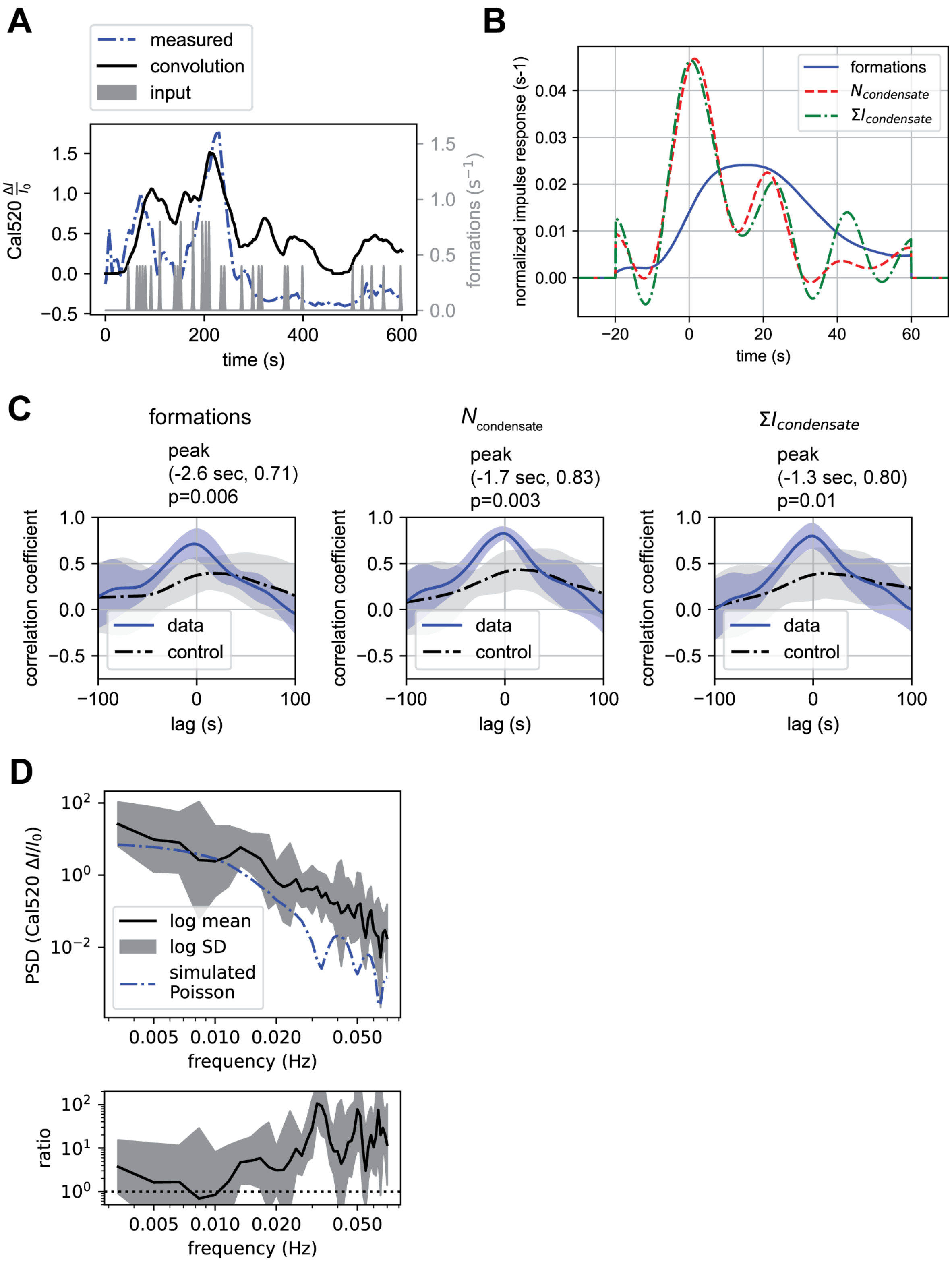
A
measured
convolution
input
Cal520 ΔI/I0
formations (s−1)
time (s)
B
formations
Ncondensate
ΣIcondensate
normalized impulse response (s-1)
time (s)
C
formations
peak
(-2.6 sec, 0.71)
p=0.006
Ncondensate
peak
(-1.7 sec, 0.83)
p=0.003
ΣIcondensate
peak
(-1.3 sec, 0.80)
p=0.01
correlation coefficient
data
control
lag (s)
D
PSD (Cal520 ΔI/I0)
log mean
log SD
simulated
Poisson
frequency (Hz)
ratio
frequency (Hz)

**Fig. S18. Spectral analyses for the dataset with higher pMHC density.**

Spectral analyses were performed for the dataset with pMHC density 1.1 ± 0.2 $\mu m^{-2}$. (**A**) Experimental condensate formation events (input), calcium trace (output), and the convolution-regenerated calcium trace from a representative cell. Convolution successfully recapitulated fluctuation pattern to a visible degree. (**B**) Comparison of the impulse responses for different inputs. Fine fluctuations appeared due to limited time resolution. (**C**) Cross-correlation between experimental calcium traces and convolution-regenerated calcium traces. False-correlation controls were calculated by comparing two traces from different cells. SD is shown as shaded regions. (**D**) PSD of calcium traces (logarithmic mean ± SD) in comparison with simulated Poisson noise (top). The ratio of experimental PSD and simulated PSD is also shown (bottom).

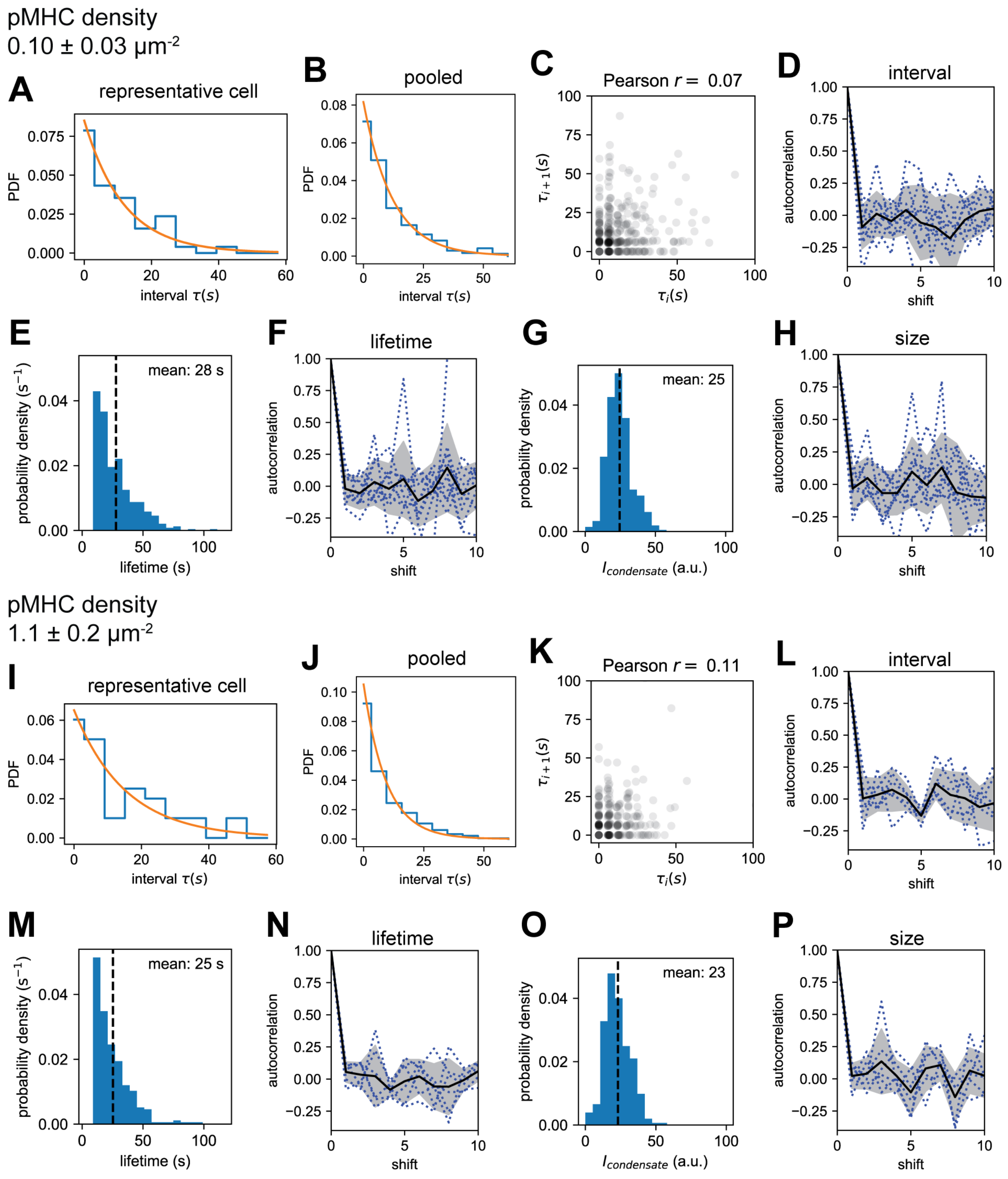


**Fig. S19. Distribution of condensate characteristics.**

(**A, B**) The histogram of intervals between condensate formation events from a representative cell or pooled from all cells. Least-square fit by exponential distribution is overlaid. (**C**) The correlation of consecutive two interval values are visualized by the scatter plot of paired values. No positive correlation was found. (**D**) Autocorrelation (defined as Pearson's correlation coefficient) of interval value sequence. Autocorrelation was calculated for each cell (blue dotted lines) and mean and SD are shown (black solid line and grey shades). Shift of one is equivalent to the correlation in (**C**), and this autocorrelation provides a generalization to larger shifts. (**E**) Distribution of condensate lifetime, pooled from all cells. (**F**) Autocorrelation of lifetime sequence. (**G**) Distribution of condensate size ($I_{condensate}$), pooled from all cells. (**H**) Autocorrelation of size sequence. (**A-H**) are for the dataset with lower pMHC densities (0.10 ± 0.03 $\mu m^{-2}$). (**I-P**) depicts the identical analyses performed for the dataset with higher pMHC densities (1.1 ± 0.2 $\mu m^{-2}$).

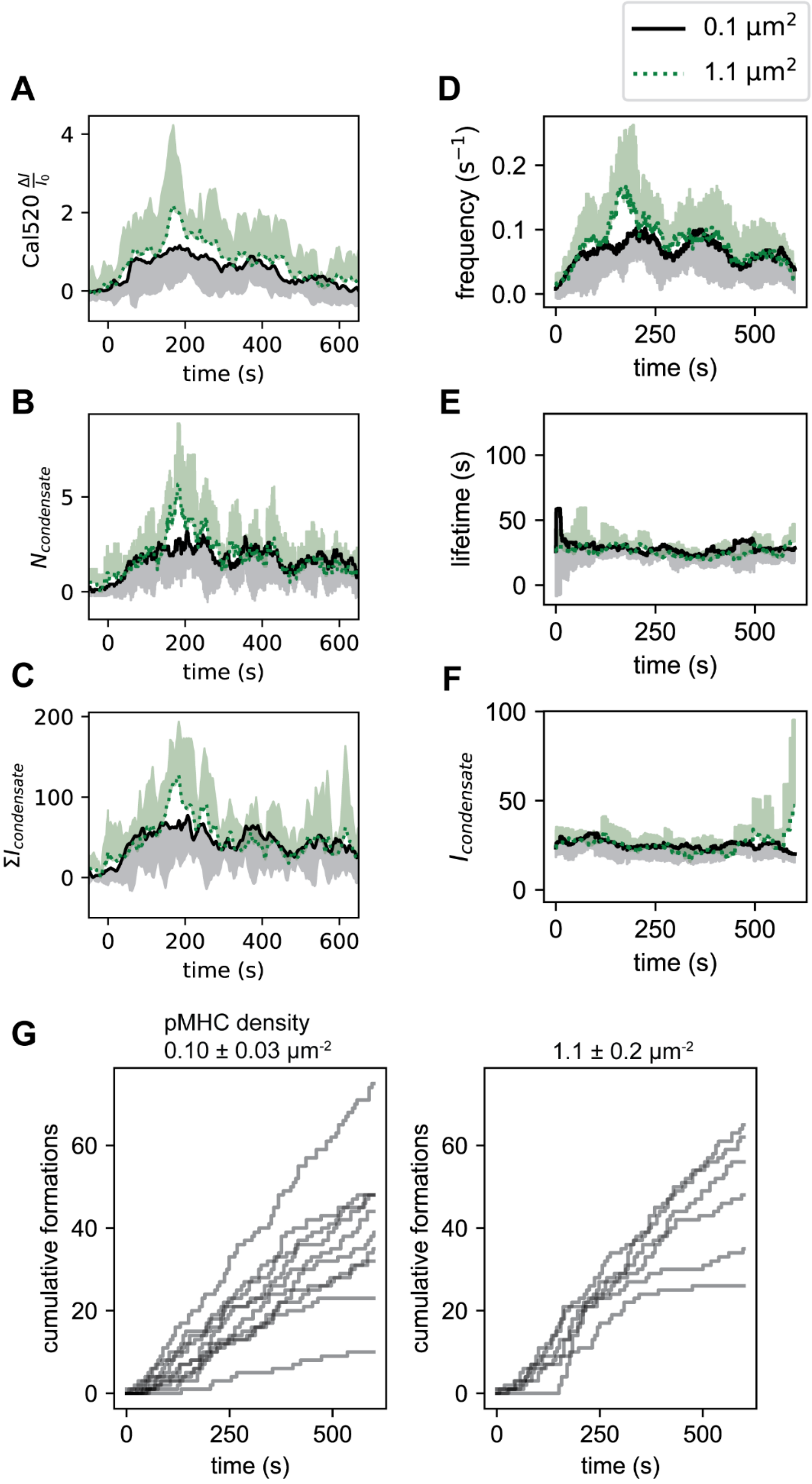
0.1 μm2
1.1 μm2
A
Cal520 ΔI/I0
time (s)
B
Ncondensate
time (s)
C
ΣIcondensate
time (s)
D
frequency (s−1)
time (s)
E
lifetime (s)
time (s)
F
Icondensate
time (s)
G
pMHC density
0.10 ± 0.03 μm-2
1.1 ± 0.2 μm-2
cumulative formations
time (s)
cumulative formations
time (s)

**Fig. S20. Uniform characteristics and non-uniform formation frequency of LAT condensates**

(**A-F**) The population-averaged $Ca^{2+}$ signal and LAT condensate characteristics for two lower and higher pMHC density. $Ca^{2+}$ signal, the momentary number of condensates, summed condensate size, and condensate formation rate exhibited a peak with higher pMHC density at about 200 sec (**A**-**D**), while the lifetime and intensity of each condensate were constant over time (**E**, **F**). The formation frequency, lifetime, and condensate intensity (frame-averaged within each condensate) were 50 s window-averaged for each cell, then population-averaged. Mean and SD are shown as half-shaded regions. (**G**) Individual cumulative formation events traces for each density range. Low density condition is the same plot as **Fig. 7A** shown for comparison (left).

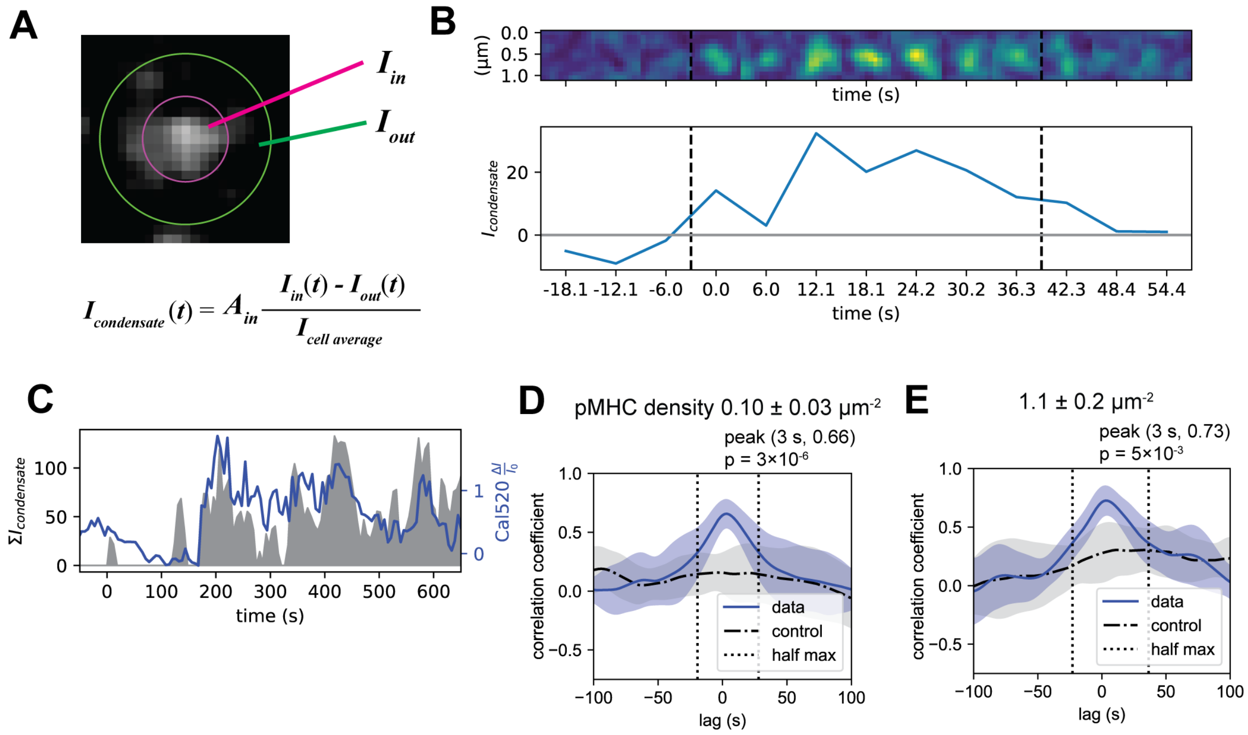


**Fig. S21. Cross-correlation of summed condensate size and calcium signal.**

(**A**) Example condensate image and calculation of condensate size $I_{condensate}$ (**B**) Example montage of a condensate and $I_{condensate}$ trace. (**C**) Example cell traces of Cal520 readout (blue line) and the summed condensate size $\Sigma I_{condensate}$ (grey area), showing clear correlations. (**D, E**) Cross-correlation of $\Sigma I_{condensate}$ and Cal520 readout at lower and higher pMHC densities.

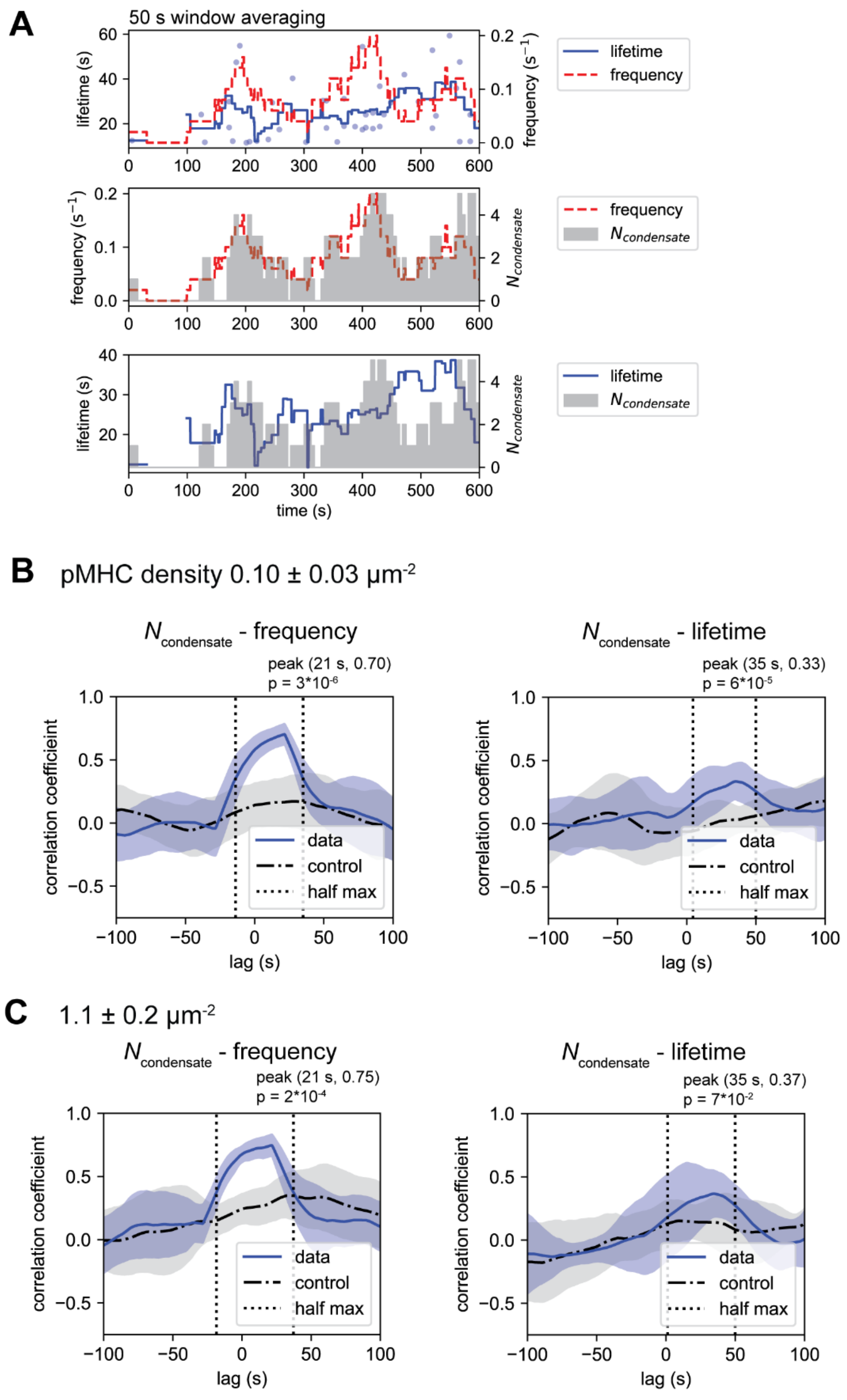
A
50 s window averaging
lifetime (s)
frequency (s−1)
lifetime
frequency
Ncondensate
time (s)
B
pMHC density 0.10 ± 0.03 µm-2
Ncondensate - frequency
peak (21 s, 0.70)
p = 3*10-6
Ncondensate - lifetime
peak (35 s, 0.33)
p = 6*10-5
correlation coefficieint
data
control
half max
lag (s)
C
1.1 ± 0.2 µm-2
Ncondensate - frequency
peak (21 s, 0.75)
p = 2*10-4
Ncondensate - lifetime
peak (35 s, 0.37)
p = 7*10-2

**Fig. S22. The momentary number of LAT condensates is majorly influenced by formation frequency.**

(**A**) Local-time formation frequency and condensate lifetime were quantified as 50 s window averages. Each lifetime and formation timing were plotted, which was then window-averaged (bottom). The local formation frequency showed more similarity to $N_{\text{condensate}}$ than local lifetime (upper and middle panels). (**B, C**) Cross-correlation of $N_{\text{condensate}}$ and each local parameter. The local formation frequency showed stronger correlation than lifetime. p value: two-tailed Welch's t-test.

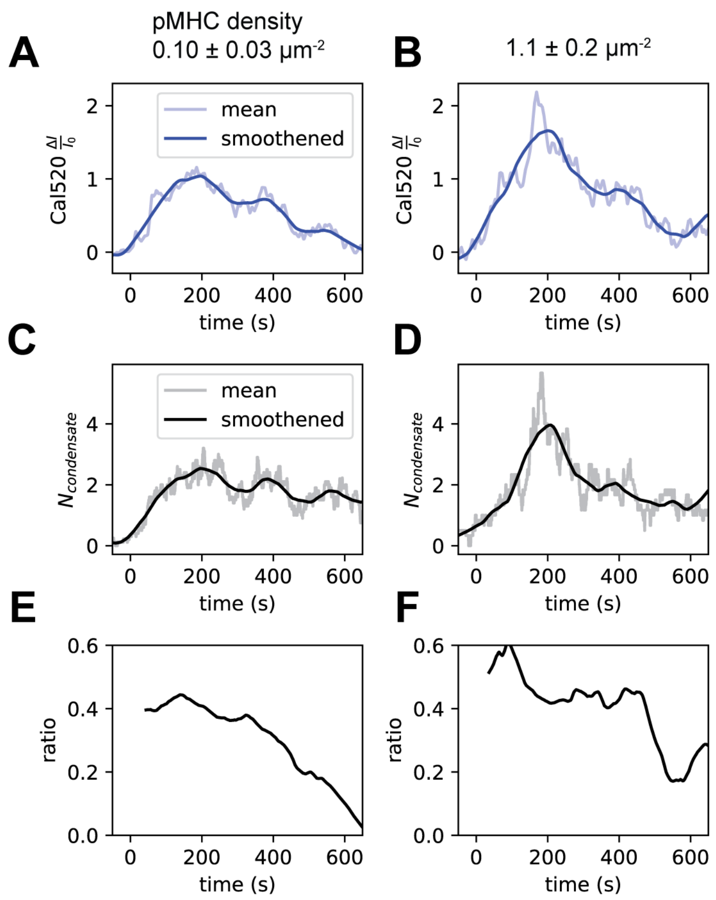


**Fig. S23. Global desensitization of calcium response to LAT condensates**

(**A, B**) The population-averaged $N_{\text{condensate}}$ and its smoothened trend (Savitzky-Golay filter) for each pMHC density. (**C, D**) The population-averaged Cal520 $\Delta I/I_0$ and its smoothened trend. (**E, F**) The smoothend Cal520 $\Delta I/I_0$ divided by the smoothened $N_{\text{condensate}}$. Slowly decreasing ratio traces indicated global desensitization of condensate-$Ca^{2+}$ response.

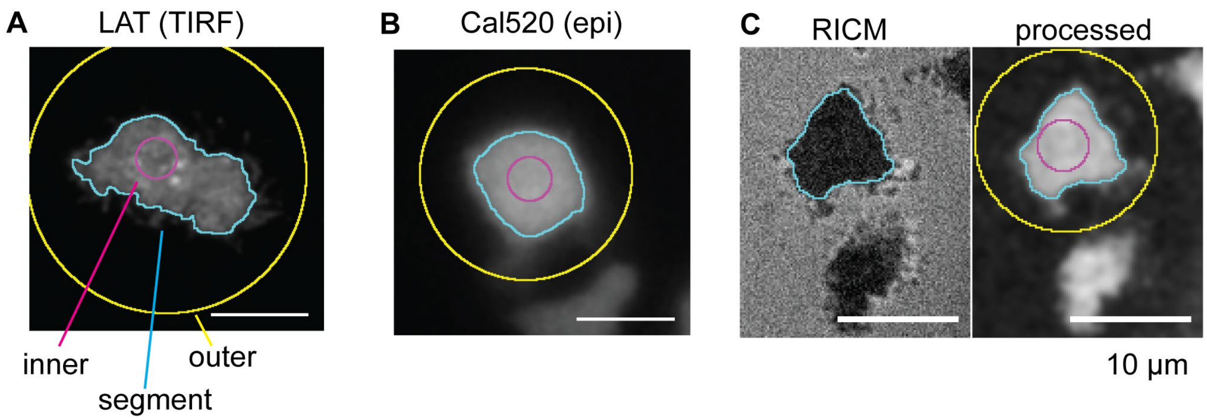


**Fig. S24. Representative images of cell segmentation by frame-by-frame thresholding**
(**A**) LAT TIRF image. (**B**) Cal520 epi-fluorescence image (**C**) RICM image of cell-foot print. RICM image was processed to produce intensity image and then segmented.

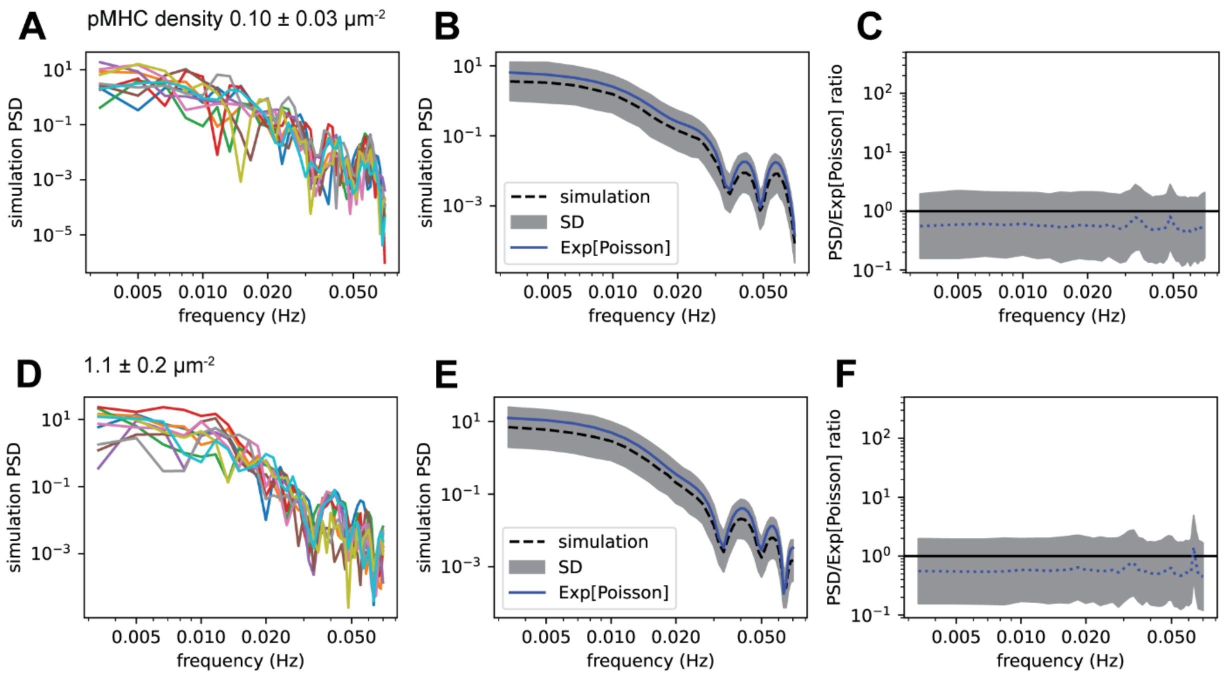


**Fig. S25. Simulation of Poisson noise and comparison with analytical approximation.** For each condition with pMHC densities of 0.10 ± 0.03 (**A-C**) and 1.1 ± 0.2 $\mu m^{-2}$ (**D-F**), condensate formation events were simulated as a Poisson process with rate 0.07 $s^{-1}$, and calcium signal trace was generated by convoluting the empirical impulse responses. These simulated calcium traces were converted to PSD in the identical way as the experimental PSD. 1000 independent traces were simulated. (**A, D**) 10 randomly chosen representative PSD traces. (**B, E**) Logarithmic mean and SD for simulated PSD. The analytical approximation of expected PSD is overlaid (blue solid line). (**C, F**) The ratio of PSD and analytical approximation shown as logarithmic mean and SD (blue dotted line and grey shade). The ratio only slightly differs from 1 (black line).

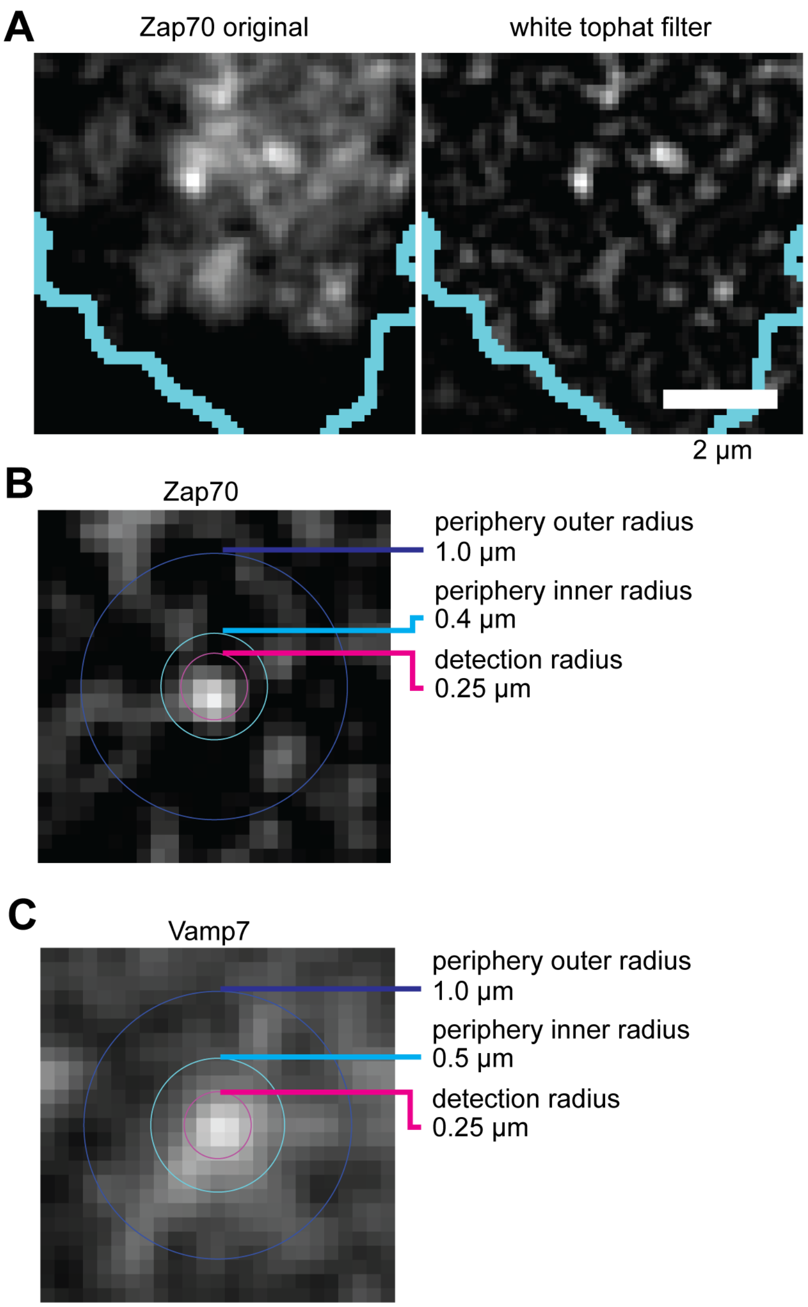


**Fig. S26. Example images for the detection of Zap70 clusters and Vamp7-associated vesicles.**

(**A**) Zap70 image was processed with white tophat filter to remove the cytosolic background signal. (**B**) Zap70 cluster with three overlaid radii used for detection criteria (see Materials and Methods). Two periphery radii were used to calculate the local background and the detection radius was used to define the condensate proximity. (**C**) Vamp7-associated vesicle with three overlaid radii.

**Table S1.**

The parameters used for cell segmentation procedure

| Channel | $\sigma$ (pixels) | $\alpha_{low}$ | $\alpha_{high}$ | $K_{opening}$ (pixels) | $A_{min}$ (pixels) |
|---|---|---|---|---|---|
| LAT (1.06 μm/pixel) | 2 | 0.5 | 1.5 | 5×5 | 400 |
| RICM (1.6 μm/pixel) | 2 | 0.5 | 1.5 | 3×3 | 200 |
| Cal520 (1.06 μm/pixel) | 2 | 0.7 | 1.5 | 5×5 | 400 |
| Cal520 (1.6 μm/pixel) | 1.5 | 0.7 | 1.5 | 3×3 | 200 |
| DiD (1.06 μm/pixel) | 2 | 0.5 | 1.5 | 5×5 | 400 |

**Movie S1.**

Imaging of pMHC:TCR (left panel), LAT condensates (middle panel), and Cal520 (right panel) of a representative cell with the pMHC density of 0.10 ± 0.03 molecules μm$^{-2}$. Time stamps indicate the time after adhesion as minutes:seconds. Scalebars indicate 10 μm.
